# Spatial population genetics in heavily managed species: Separating patterns of historical translocation from contemporary gene flow in white-tailed deer

**DOI:** 10.1101/2020.09.22.308825

**Authors:** Tyler K. Chafin, Zachery D. Zbinden, Marlis R. Douglas, Bradley T. Martin, Christopher R. Middaugh, M. Cory Gray, Jennifer R. Ballard, Michael E. Douglas

**Author notes:** Email: (TKC); (BTM); (ZDZ); (MED); (MRD), Email: (CRM); (JRB); (MCG).

## Abstract

Approximately 100 years ago, unregulated harvest nearly eliminated white-tailed deer (*Odocoileus virginianus*) from eastern North America, which subsequently served to catalyze wildlife management as a national priority. An extensive stock-replenishment effort soon followed, with deer broadly translocated among states as a means of re-establishment. However, an unintended consequence was that natural patterns of gene flow became obscured and pre-translocation signatures of population structure were replaced. We applied cutting-edge molecular and biogeographic tools to disentangle genetic signatures of historical management from those reflecting spatially heterogeneous dispersal by evaluating 35,099 single nucleotide polymorphisms (SNPs) derived via reduced-representation genomic sequencing from 1,143 deer sampled state-wide in Arkansas. We then employed Simpson’s diversity index to summarize ancestry assignments and visualize spatial genetic transitions. Using sub-sampled transects across these transitions, we tested clinal patterns across loci against theoretical expectations of their response under scenarios of recolonization and restricted dispersal. Two salient results emerged: (A) Genetic signatures from historic translocations are demonstrably apparent; and (B) Geographic filters (major rivers; urban centers; highways) now act as inflection points for the distribution of this contemporary ancestry. These results yielded a state-wide assessment of contemporary population structure in deer as driven by historic translocations as well as ongoing processes. In addition, the analytical framework employed herein to effectively decipher extant/historic drivers of deer distribution in Arkansas are also applicable for other biodiversity elements with similarly complex demographic histories.

## 1 INTRODUCTION

Understanding movement behaviors and patterns of dispersal across the landscape are important components of a wildlife management strategy. Quantifying dispersal and connectivity among populations has traditionally been done using direct methods such as radio- and satellite-tracking (Kays, Crofoot, Jetz, & Wikelski, 2015), yet these methods are labor-intensive, generally only allow for small sample-sizes with results difficult to extrapolate at the population level (Katzner & Arlettaz, 2020). An ability to do so indirectly via patterns measured from genetic data such as gene flow (i.e., corresponding to migration, demographics) and genetic drift (i.e., effective population sizes, inbreeding) has made it possible to more accurately assess movement and landscape ecology patterns at a fraction of the per-individual cost (Bossart & Pashley Prowell, 1998; Comte & Olden, 2018; Lowe & Allendorf, 2010). Continual decline of sequencing costs concomitant with computational and analytical advancements now offer an unprecedented density of sampling and geographic resolution that can link individual movement patterns with landscape genetic associations (Richardson, Brady, Wang, & Spear, 2016), as well as functional genomic correlations (Schoville et al., 2012). This has spurred a proliferation of approaches, such as deducing natal assignment (Battey, Ralph, & Kern, 2020a), inferring the directionality and pathways of range expansions (Rengefors et al., 2020), and predicting disease spread from spatial host-pathogen associations (Fountain-Jones et al., 2021; Kozakiewicz et al., 2018).

However, contemporary evaluations examine but a single ‘snapshot’ from an evolutionary continuum of events that have cumulatively shaped contemporary diversity, a consequence of which is that genetic patterns may not be interpretable when juxtaposed against modern landscapes (Douglas, Minckley, & DeMarais, 1999). Thus, models that invoke landscape genetic associations may incorporate implicit assumptions that lack biological reality (Cushman & Landguth, 2010). For example, one common assumption involves dispersal among a series of discontinuous sub-populations (Lundgren & Ralph, 2019; Petkova, Novembre, & Stephens, 2015) that may incorrectly infer discrete causes (such as barrier effects or reproductive isolation)(Bradburd, Coop, & Ralph, 2018). Likewise, gene flow and genetic drift are often conflated, in that one may generate spurious signals of the other (Battey, Ralph, & Kern, 2020b; Mazet, Rodríguez, Grusea, Boitard, & Chikhi, 2016). For example, demographies with a history of turbulence may inflate genetic divergences such that population structure appears artificially discrete and thus suggests restricted gene flow (Austerlitz, Jung-Muller, Godelle, & Gouyon, 1997; Excoffier & Ray, 2008; Nei, Maruyama, & Chakraborty, 2014). A classic example of this phenomenon is the re-expansion of populations following an historical ebb. Upon secondary contact, unique genetic changes that have accumulated during periods of isolation may seemingly initiate an apparent rapid shift in ancestries when viewed from a landscape genetic context (Zellmer & Knowles, 2009).

More recent events such as anthropogenic habitat fragmentation may also act to obscure genetic patterns across the landscape (Epps, Wasser, Keim, Mutayoba, & Brashares, 2013; Landguth et al., 2010; Pavlacky, Goldizen, Prentis, Nicholls, & Lowe, 2009). In particular, human-mediated dispersal introduces both diversifying and homogenizing effects that prove difficult to disentangle (Epps & Keyghobadi, 2015; Seebens et al., 2017). This erodes adaptive and/ or spatial associations in that it disturbs the expected relationships between genetic differentiation and geographical or environmental proximity (Capinha, Essl, Seebens, Moser, & Pereira, 2015; Einfeldt, Jesson, & Addison, 2020). Well-intended actions, such as translocations and stockings, may also inadvertently obscure pre-existing landscape genetic patterns (Brown, Hull, Updike, Fain, & Ernest, 2009; Laikre, Schwartz, Waples, & Ryman, 2010; Shephard, Ogden, Tryjanowski, Olsson, & Galbusera, 2013). This is especially so for heavily-managed species that incur translocations as one aspect of their management (Jahner et al., 2019; Seddon, Griffiths, Soorae, & Armstrong, 2014)

White-tailed deer, one of the most recreationally important species in North America, is a case in point (Knoche & Lupi, 2012), with populations intensely impacted by both hunter harvest and game management (Waller & Alverson, 1997; Wolverton, Kennedy, & Cornelius, 2007).

Early in the 20^th^ century, deer and other species extensively hunted (e.g., wild turkey) saw widespread extirpations. In a census of the north-central United States, Leopold (1931) noted that extant deer populations were of low abundance in southern Missouri, an observation that dovetails with estimated numbers of deer in Arkansas during that period (∼500 scattered across isolated refugia at the low ebb; Holder, 1951). Subsequent efforts to bolster resident numbers and re-populate depleted areas in Arkansas involved translocations from within- and out-of-state. Although successful, these efforts promoted genetic patterns within the state that were based on artificial, rather than natural, movements.

Distributional patterns in white-tailed deer clearly reflect multiple factors. Those extrinsic may include: Rivers, interstate highways, suitable habitat (Robinson, Samuel, Lopez, & Shelton, 2012); agricultural land use and urbanization (Kelly et al., 2014); as well as regional eco-physiographic boundaries (Miller, Miller-Butterworth, Diefenbach, & Walter, 2020). Intrinsic factors, on the other hand, may point to: Population density (Lutz, Diefenbach, & Rosenberry, 2015); age structure/ sex ratio (Long, Diefenbach, Rosenberry, & Wallingford, 2008); and social hierarchy (Nixon & Mankin, 2016).

A first step in designating factors that have shaped deer population structure and dispersal is to estimate the genetic component within Arkansas deer genomes that still persists as a residual of historic translocations and subsequent genetic drift. Perhaps our most pressing need with regard to dispersal is the impact these data may have in relation to a widespread and fatal neurodegenerative disease of cervids (chronic wasting disease; CWD), which now represents a panzootic (Williams and Young 1980; Escobar et al. 2019; Mawdsley 2020). Thus, efforts to contain and mitigate its spread are paramount for wildlife management, not only in North America, but also globally (Leiss et al., 2017).

### 1.1 Separating contemporary dispersal from historical translocation

The driving factors for deer dispersal primarily relate to habitat quality and climate, and these in turn elicit obvious management interest. Region-specific patterns are needed so as to develop broader generalizations regarding species-specific movement ecology (Brinkman, Deperno, Jenks, Haroldson, & Osborn, 2005), and to evoke management strategies that can optimize hunter-harvest while maintaining population densities.

The goal of our study was to ascertain if signatures of historic translocations are apparent in, and have contributed to, the genetic diversity and structure of deer in Arkansas, and furthermore, if they can indeed be parsed from a confusing matrix of ongoing gene flow and genetic drift. To do so, we surveyed a broad array of nuclear genomic markers using ddRAD sequencing (Peterson et al., 2012). Our first objective was to characterize respective patterns of anthropogenic translocation and natural re-colonization via refugial populations, and secondarily to seek evidence for geographic barriers throughout the state that seemingly prevent or actively filter deer dispersal. However, we hypothesized that the former situation would obscure the latter, in that long-range anthropogenically-mediated displacement of individuals violates methodological assumptions that stipulate gene flow as occurring in a spatially consistent manner (Bradburd & Ralph, 2019).

Here, we used ancestry assignment probabilities (e.g., for *K*-number of populations) and reduced this to a single index (Simpson, 1949) that encapsulated the diversity of ancestries within a given spatial grain. This served to reduce those genetic ancestry components that are influenced by widespread translocation, to a single interpolated ‘surface’ representing only changes in ancestry over space (i.e., boundaries between contemporary populations). We also wished to define the respective roles played by stochastic and deterministic processes in generating these zones, particularly given historic population fluctuations and the internal/ external translocations that conflated the demography of Arkansas deer. To do so, we borrowed components of cline theory to hypothesize how individual loci should transition across these spaces, relative to a genome-wide average (Barton, 1983; Barton & Hewitt, 1985; Endler, 1973; Hewitt, 2001; Polechová & Barton, 2011; Slatkin, 1973).

Although clinal variation is often taken as evidence for selection (i.e., migration-selection balance; Haldane, 1948), genetic drift in concert with spatially variable gene flow can generate patterns in individual loci that mimic those seen in adaptive clines (Vasemägi, 2006). With drift operating alone, allele frequencies at each locus may wander ‘upward’ or ‘downward’ across space as a function of the initial allele frequency. Range expansion or dissemination from refugia can also yield a type of ‘rolling’ founder effect that can mirror a clinal pattern (Excoffier, Foll, & Petit, 2009; Hallatschek & Nelson, 2008; Hewitt, 2000; Keller et al., 2013). Accordingly, we might predict that multi-locus patterns at population boundaries generated solely due to genetic drift in expanding populations will be dominated by stochastic directional change, with each fluctuating either above or below the genome-wide average (Santangelo, Johnson, & Ness, 2018).

On the other hand, a strong barrier to movement would yield a discontinuity in the rate of directional change across space (Barton, 2008; Nagylaki, 1976; Slatkin, 1973). The existence of a barrier (either impacting dispersal or as a strong fitness differential) thus implies the presence of an inflection point that would induce a rapid shift in locus-specific ancestries (the ‘width’ of this cline is the inverse of the slope at the inflection point; Slatkin 1973; Endler 1977; Fitzpatrick 2013).

Given these predictions, we summarized locus-wise patterns using two clinal parameters: *α*, which describes the direction of genetic change, and *β*, which indicates ‘width’ of a cline. We examined how these two parameters varied, then developed hypotheses regarding the manner by which it impacted our observed population structure. We then considered the evolutionary implications, and limitations, of these results in the context of management, and in so doing demonstrated how the methodologies employed herein can be used to make inferences in other heavily managed species regarding demographic factors (i.e., sex-biased and natal dispersal) and those more extrinsic (i.e., landscape resistance) that drive population structure.

## 2 METHODS

### 2.1 Sampling and data collection

During 2016-2019, 1,720 tissues were collected by the Arkansas Game and Fish Commission (AGFC), representing all 75 Arkansas counties. We employed a combination of targeted sampling, road-kill surveys, and a voluntary state-wide CWD testing program (Chafin et al., 2020). Age and sex were collected where possible, with the former estimated by tooth development and wear (Severinghaus, 1949). Data for an additional 30 samples were also attained from Wisconsin to test for signals of historically recorded translocation efforts involving this stock (Holder, 1951). From these, a subset of 1,208 samples were chosen for sequencing.

We homogenized tongue or ear tissue (stored at −20°C) and extracted genomic DNA using QIAamp Fast Tissue kits (Qiagen, Inc), with verification via gel electrophoresis (2% agarose). Samples with sufficient yields of high molecular weight DNA (>200ng) were then enzymatically fragmented via incubation at 37°C, using high-fidelity *NsiI* and *MspI* restriction enzymes (New England Biolabs, Inc.), following enzyme and size-selection optimization using *in silico* digests (Chafin, Martin, Mussmann, Douglas, & Douglas, 2018) of several available reference genomes hosted by NCBI: *Odocoileus virginianus* (GCA_002102435.1), *Capreolus capreolus* (GCA_000751575.1), and *Capra hircus* (GCF_001704415.1).

Digests were purified using Ampure XP beads (Beckman-Coulter, Inc.) and standardized to 100ng per sample. Unique inline barcodes (Peterson et al., 2012) were then ligated using T4 DNA Ligase (following manufacturer protocols; New England Biolabs, Inc.). Samples were then multiplexed (N=48) prior to automated size selection at 300-450bp using a Pippin Prep (Sage Sciences). Adapter extension was performed over 12 PCR cycles using TruSeq-compatible indexed primers (Illumina, Inc.) and Phusion high-fidelity *taq* polymerase (New England Biolabs, Inc.) Additional quality controls (e.g., qPCR and fragment analysis) were performed on final libraries prior to 1×100 single-end sequencing on the Illumina HiSeq 4000 (Genomics and Cell Characterization Facility, University of Oregon/Eugene), with a total of N=96 samples pooled per lane.

Raw reads were demultiplexed using the pyRAD pipeline (Eaton, 2014), and those with barcode mismatches were discarded. Demultiplexed reads were further filtered by removing those having >4 nucleotides below a quality threshold of 99% accuracy. Reads were then clustered into putative loci within-individuals, allowing for a maximum distance threshold of 15%. This was done using the VSEARCH algorithm (Rognes, Flouri, Nichols, Quince, & Mahé, 2016) as implemented in pyRAD, so as to remove read clusters with 3+ indels, >5 ambiguous consensus nucleotides, or a coverage <20X or >500X. Putative homologs were identified using among-individual clustering with the same parameters, and additional removal of loci having >2 alleles per individual, >70% heterozygosity for any polymorphic site, >10 heterozygous sites, or <50% individual recovery (see *github*.*com/tkchafin/scripts* for post-processing and file formatting scripts). To mitigate issues of independence among sites, we additionally sub-sampled the dataset to one SNP per locus.

### 2.2 Derivation of population structure

Given known issues with respect to assignment accuracy in datasets dominated by low-frequency variants (Linck & Battey, 2019), we excluded SNPs exhibiting a minor allele count <2 which corresponds to a global minor-allele frequency of ∼1%. This minor-allele count threshold per-locus (i.e., accounting for total number of sampled alleles), equates to a mean minor-allele frequency threshold of 11% (sd=2%). We then inferred population structure (ADMIXTURE: Alexander et al., 2009) with parallel processing (ADMIXPIPE: Mussmann et al., 2020). Model selection (i.e. for *K*, the number of populations) followed a cross-validation approach with results aggregated from 20 independent replicates (CLUMPAK: Kopelman et al., 2015).

Individual-level ADMIXTURE results were summarized as a ‘surface’ with spatial discontinuities represented as interpolated assignment probabilities. Here, we constructed state-wide rasters, as representing per-pixel probabilities or ‘ancestry proportions,’ using Empirical Bayesian Kriging (ARCMAP 10.7.1, Esri, Inc.). Probability surfaces were then summarized as evenness and diversity of ancestries in each cell using Simpson’s index (Simpson, 1949) (where *K*=number of statewide sub-populations). Our use of the diversity index was based on a straightforward prediction: Areas representing spatial transitions between populations will have a correspondingly low certainty of assignment to a given sub-population ancestry (i.e., high diversity). Likewise, those with low inter-population exchange will comprise genetically similar individuals assigned with high probability to the endemic ancestry (i.e., low diversity). Thus, genetic edges represent a marked transition from one population to another, as identified by site-wise diversity in assignment probabilities (i.e. ‘ancestry diversity’). By lowering the resolution of the genetic diversity surface, we effectively decreased the noise introduced by translocation.

We expect spatial variation in ancestry diversity to be inversely proportional to true rates of gene flow, in that this quantity (as well as the ancestry proportions from which it is computed) are a product of gene flow averaged over many generations. Of note is the fact that the method only examines local ancestry probabilities and consequently will not be dominated by translocation-related artefacts (as are methods based on interpolated pairwise genetic distances or *F*_ST_). Patterns based on Simpson’s diversity were contrasted with those inferred using a form of 2D-stepping stone model (EEMS; Petkova et al. 2015), as run with 2 million MCMC iterations (1 million as burn-in), and sampled every 1,000 iterations [following parameter sweeps that tuned MCMC acceptance rates to fall between 20% and 50% (Roberts, Gelman, & Gilks, 1997)].

### 2.3 Estimating cline parameters

We examined the nature of population boundaries by examining how individual loci transitioned across these regions, relative to the genome-wide average. To do so, we defined eight transects across putative genetic edges, sampling 32-73 individuals per transect 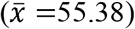. Individuals were chosen to represent each sub-population using a probability threshold applied to ADMIXTURE results. Loci were filtered to remove all SNPs with missing data in >50% of individuals, and computational time was further restricted by retaining only those loci with a sufficient allele frequency differential among clinal extremes [computed as δ > 0.50; (Gregorius & Roberds, 1986)]. Locus-wise clinal patterns were then inferred using a Bayesian method developed originally for hybrid zones (BGC: Gompert & Buerkle, 2011, 2012). Open-source Python code for filtering SNP matrices and generating necessary input are available at: (*github*.*com/tkchafin/scripts/phylip2introgress*.*pl* and *phylip2bgc*.*pl*).

Analyses were performed for each transect across four replicates, each using one million MCMC iterations, discarding the first 500,000 as burn-in, with output thinned to every 500 iterations. Results were summarized by visualizing a 2-D density of cline shape parameters. These are: α (=cline directionality) that describes an increase (α > 0) or decrease (α < 0) in the probability of locus-specific ancestry from a parental population; and β (=cline width/ steepness) that defines the rate of transition in probabilities of locus-specific ancestries having either steep (β > 0) or wide (β < 0) shapes (Gompert, Parchman, & Buerkle, 2012). In this context, a locus which does not deviate from the genome-wide pattern would have α=β=0. Deviation in a directional manner (i.e., an increase or decrease of one ancestry over another) is described by α, whereas deviation in the rate of ancestry change around an inflection point (i.e., sigmoidal) is described by β. Statistical outliers were designated using the method of Gompert and Buerkle (2011). BGC results were parsed and visualized using the ClinePlotR R package (Martin, Chafin, Douglas, & Douglas, 2020).

### 2.4 Relative dispersal by age and sex

Of particular interest in wildlife management is the backwards inference of geographic positioning from genotypes—that is, the geolocation of ‘origination’ points for sampled animals. This could be used, for example, to ascertain the geographic origin of poached individuals, or to estimate post-natal dispersal. To this end, we used the novel, deep-learning method LOCATOR (Battey et al., 2020a) to predict the geographic origin of samples without relying upon explicit assumptions about population genetic processes underlying spatial genetic differentiation (Bradburd & Ralph, 2019). The analysis was performed iteratively across each individual, using the remaining samples to train the LOCATOR classifier, with 100 bootstrap pseudo-replicates to assess variance in geolocation. Given computational constraints, we performed the analysis using a subset of 5000 SNPs having a minor-allele frequency >10%.

We estimated relative dispersal distances as the Euclidean distance between sampled localities and the centroid of predicted coordinates, under the assumption that the distance between predicted and collected locations is the result of lifetime dispersal, at least for samples for which geolocation variance is low among pseudo-replicates. The results were then partitioned by age, sex, and CWD-status.

Our second approach examined the decay in genetic relatedness as a function of distance from each individual, measured as a Prevosti distance (R-package poppr: Kamvar et al., 2014). Here, the assumption is that recently dispersed individuals will be, on average, more genetically dissimilar from resident individuals, whereas resident individuals having an appreciable reproductive output will be less so. These calculations were limited to individuals that had neighboring samples within a 5km radius, thus removing individuals from sparsely sampled regions where the relationship between relatedness and distance would be unreliable. We also note that the traditional aging method employed herein (Severinghaus 1949) has an accuracy seemingly reduced in older deer, potentially suggesting caution in the interpretation of results (Cook & Hart, 1979; Gee, Holman, Causey, Rossi, & Armstrong, 2002; Mitchell & Smith, 1991).

## 3 RESULTS

### 3.1 Data processing

Our raw data represented N=1,143 samples, including N=29 from Wisconsin (Table S1). We removed N=83 that had missing metadata, discrepancies with coordinates, or <50% of loci present. Assembly in pyRAD yielded an average of 25,584 loci per sample (σ=8,639). After removing loci present in <50% of samples and excluding those containing potential paralogs (e.g., excessive heterozygosity or >2 alleles per locus), our final dataset contained 35,420 loci, from which 2,655,584 SNPs were catalogued. Of these, 54,102 were excluded as singletons. To limit signal redundancy, we then condensed the data to one SNP per locus, yielded a final matrix of 35,099 SNPs for analyses of population structure.

### 3.2 Population structure and ‘ancestry surfaces’

Cross-validation, performed on N=20 replicates each for subpopulation model (*K*=1-20), revealed the optimal number of clusters as *K*=8. Spatial orientation of these samples (Fig. 1) provided a geographic definition, with some subpopulations qualitatively defined by apparent landscape features, such as the Arkansas River Valley as the southern extent of subpopulations *k*=3 and *k*=6.

**Figure 1:**
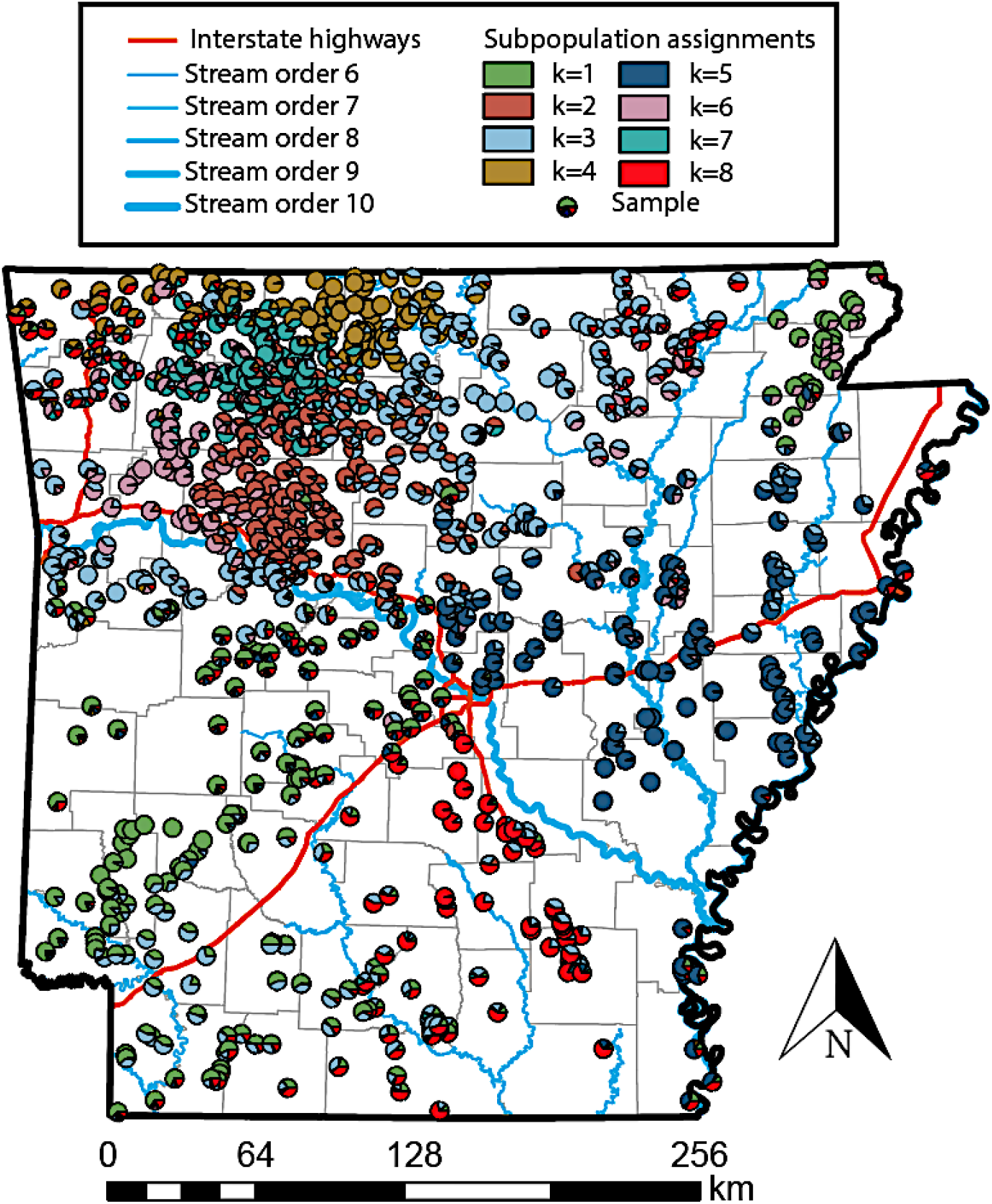
Ancestry proportions for *N*=1,143 Arkansas white-tailed deer, as inferred using the program ADMIXTURE applied to 35,099 single-nucleotide polymorphisms (SNPs) generated via ddRAD-seq. Samples are represented as pie charts plotted at absolute collection coordinates, with colors of assignment probabilities proportional to a particular subpopulation.

Two ancestries (*k*1 and *k*8) largely dominated south of the Arkansas River, bounded by Interstate 30 to the north and the Ouachita River to the south (Figs. 1-2), each of which supports the argument that genetic structure is defined by large geographic barriers. The southwestern portion of the state has two ancestral assignments (*k*1 and *k*3), with the latter having mixed representation in the north-central section (potentially an artefact of weak differentiation rather than true shared ancestry). The southeastern section is dominated by a single gene pool (*k*8), which coincidentally subsumed all Wisconsin samples, a strong signature of genetic variability as a residual of historic translocations.

A greater amount of locally endemic structure occurred north of the Arkansas River in the Ozark Mountains, where six sub-populations were evident. The most broadly distributed (*k*5) was to the east in the Mississippi alluvial plains, extending westward across the mainstem of the White River then northward towards the confluence of the Black and White rivers, where it grades into several distinct yet loosely defined sub-populations (Figs. 1-2). The northwestern corner of the state was the most heterogeneous, with four primarily endemic gene pools (*k*=2,3,4 and 7; Fig. 2). The northern-most of these was approximately bounded by the White River (Fig. 2) and graded westward into an area with high levels of mixed assignment (Fig. 1). The remaining northwestern region was defined by several gene pools showing spatially weak transitions, suggesting reduced gene flow but with geographic and/ or environmental boundaries reasonably porous.

**Figure 2:**
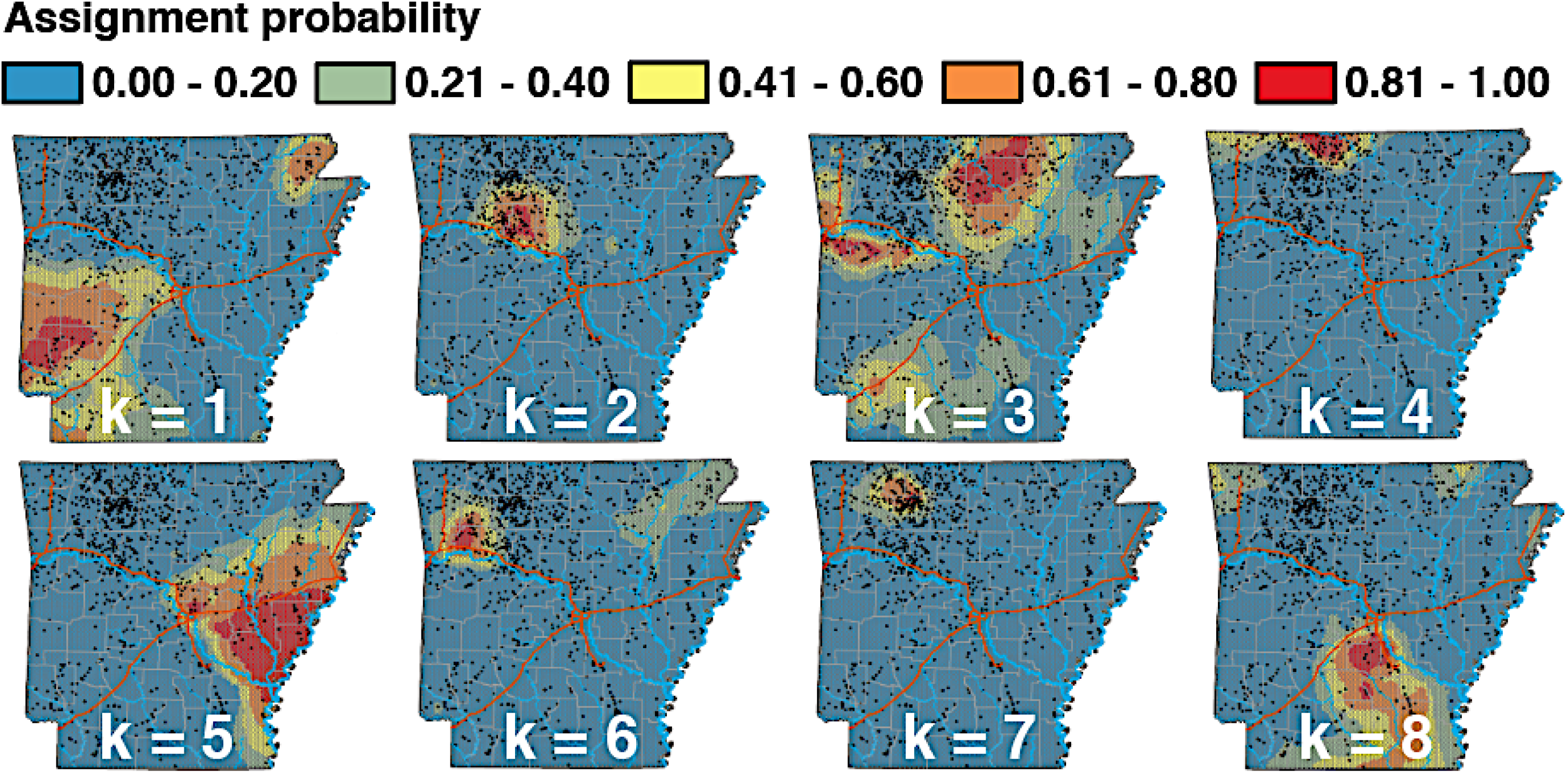
Assignment probabilities for eight populations of Arkansas white-tailed deer (*k*=1 through *k*=8), as inferred from ADMIXTURE applied to 35,099 SNPs, interpolated using Empirical Bayesian Kriging of assignments for *N*=1,143 individuals. P(*k*)=1.0 corresponds to 100% probability of ancestry per raster cell, and P(*k*)=0.0 corresponds to 0%. Individual samples are represented as black dots.

Effective migration surfaces (EEMS) failed to capture any discernable pattern relating to spatially defined population structure (Fig. S1). Geographic breaks separating sub-populations (=genetic edges) were captured instead by reducing interpolated assignment probabilities (n = *k*) as a continuous Simpson’s diversity index (n=1) (Fig. 3). This, in turn, reflects a dependence on homogeneity of local assignments, rather than global patterns compounded by long-distance transplants.

**Figure 3:**
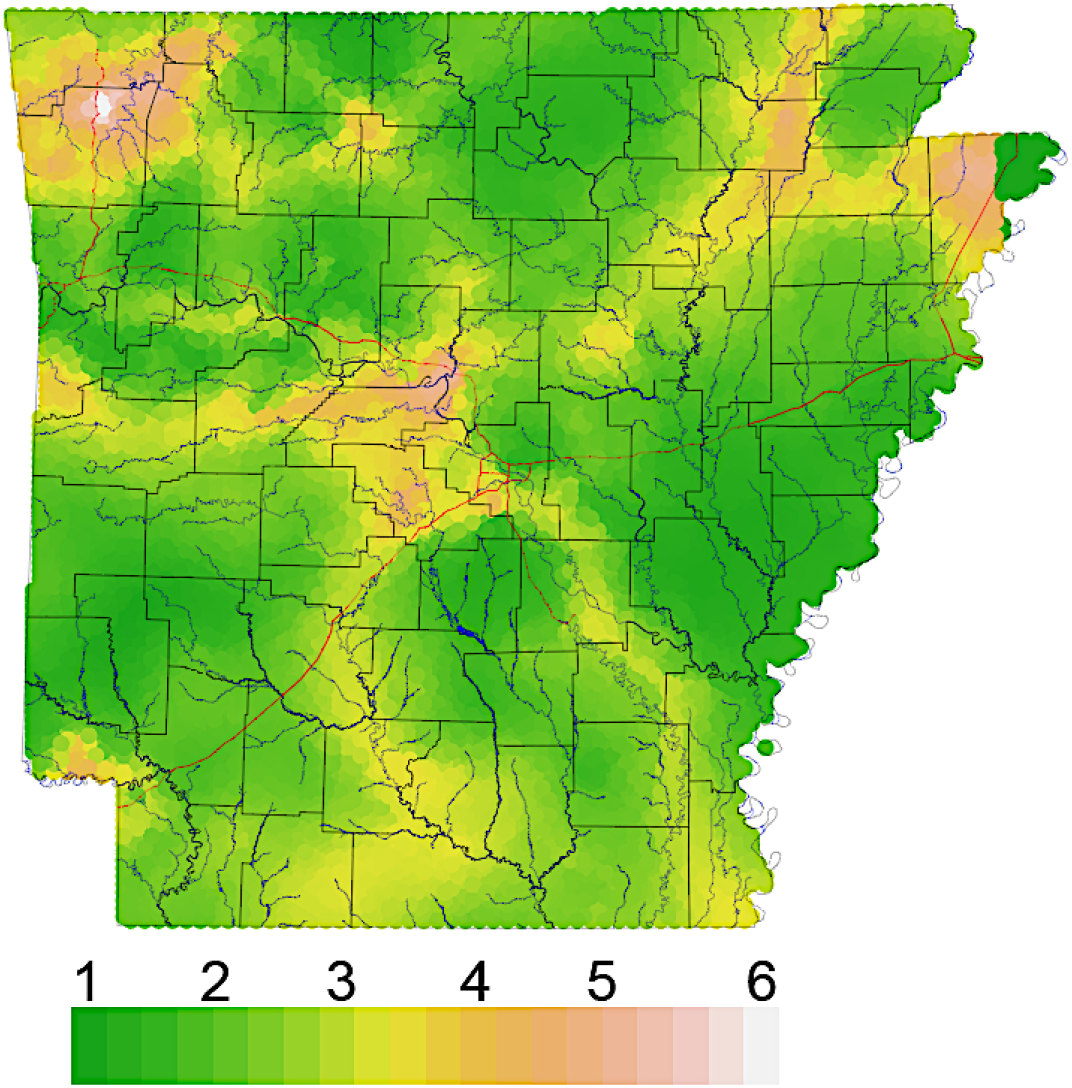
Simpson’s diversity of interpolated ancestry proportions for *N*=1,143 Arkansas white-tailed deer. Each raster cell was assigned eight values equal to the expected proportion of ancestry corresponding to the eight genetic clusters (Fig. 1), then summarized using Simpson’s diversity index.

### 3.3 Intraspecific genomic clines

Genomic clines varied substantially among transects (Figs. 4, S2). Most inter-population comparisons within northwest Arkansas, to include *k*2 x *k*3 (both eastern and southern transition zones), *k*2 x *k*6, and *k*4 x *k*7, indicated variation primarily restricted to cline directionality (α), with cline steepness (β) at a minimum. The variation in locus-wise pattern for these cases (hereafter termed ‘α-dominant’), indicated a directional change in the representation of reference populations across the transect, but without a noticeable ‘inflection’ point, as implied by non-zero cline width (β).

**Figure 4:**
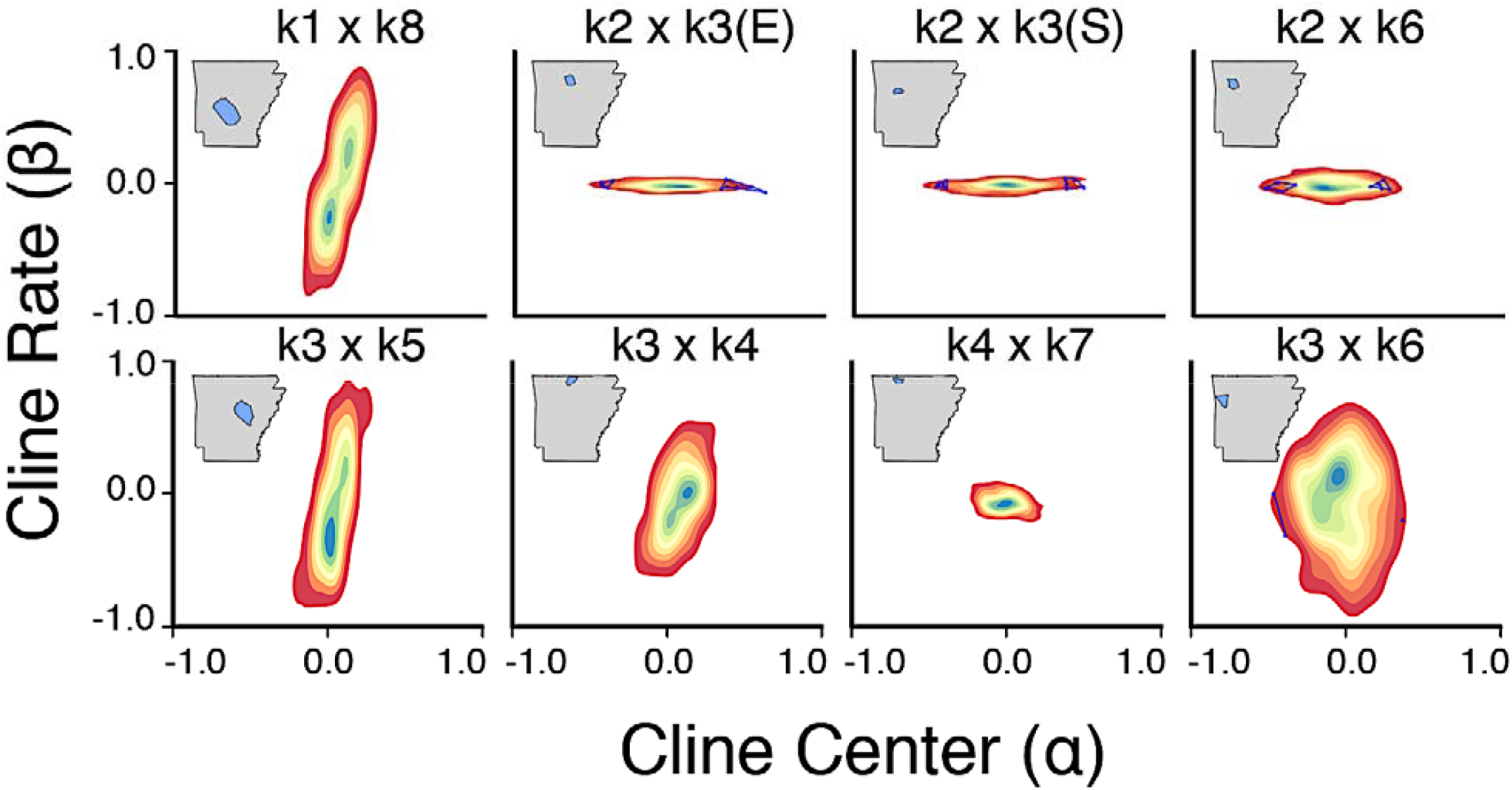
Relationships between genomic cline parameters contrasted among eight sub-sampled transects of individuals spanning population boundaries in Arkansas white-tailed deer (Fig. 3). Contour plots show relative densities of SNPs varying in cline steepness (β), representing the gradient of the clines, and cline directionality (α), representing bias in SNP ancestry. Outlier loci are highlighted in blue.

Remaining comparisons, including one additional transect from northwest Arkansas (*k*3 x *k*4), primarily reflected variation in cline width (β), in that loci varied most prominently regarding steepness of transition around an inflection point respective to genome-wide ancestry. Although two transects showed minor exception (i.e., *k*3 x *k*6 varied along both parameters; *k*4 x *k*7 each showed minimal variation), the contrasting variation between α-dominant versus β-dominant transects suggests that different processes underly ancestry transitions.

### 3.4 Estimating dispersal using geolocation analysis

Individuals from densely sampled regions could be assigned to geographic origin using the ‘deep-learning’ approach, with CWD-positive individuals assigned with a mean bootstrap distance from centroid prediction generally <15km (Fig. S3). However, we did find assignment error (e.g., among bootstraps) was elevated in low-density sampling regions (Figs. 5, S4), which resulted in higher estimated individual dispersal distances (Fig. S4A, B). Given that variance in dispersal estimates dropped considerably below ∼25km (Fig. S4C), a conservative threshold of 10km was chosen and all individuals having a mean bootstrap-centroid distance above that were removed for the purposes of dispersal estimates. After filtering, N=110 samples remained (Fig. 5A). A higher error threshold (i.e., 20km) allowed a greater number of total samples to be incorporated (N=264; Fig. 5B), with several ‘roadkills’ appearing as long-distance transfers. Although these results are low-precision assignments, they do underscore the capacity of the method regarding the identification of transported individuals (e.g., poached or illegally dumped deer, or carcasses transported across state lines or regional management zones).

**Figure 5:**
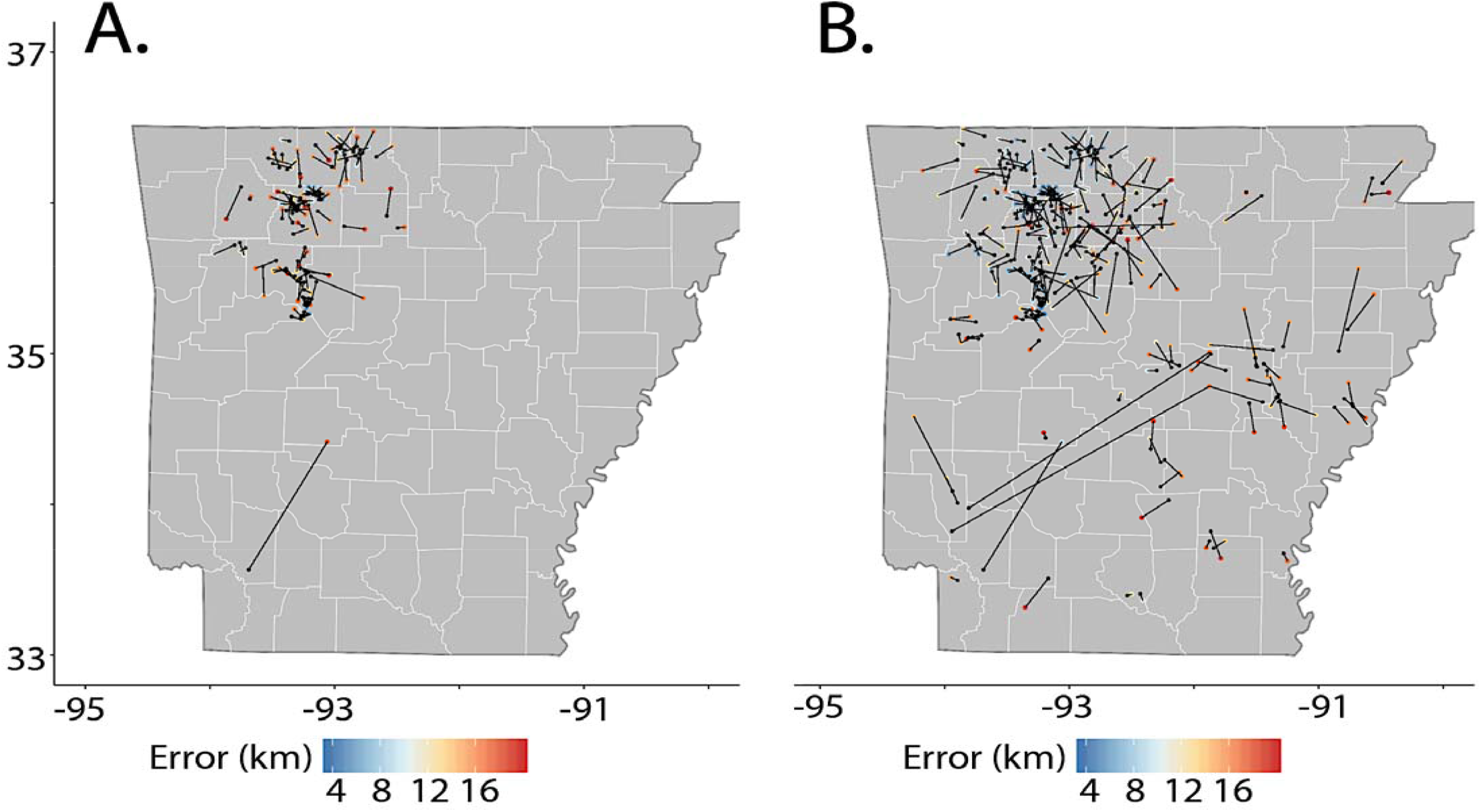
Summary of geo-location predictions in LOCATOR as a random subset of *N*=490 Arkansas white-tailed deer at two error thresholds: 10km (=A) and 20km (=B), thus constraining results to N=110 (=A) and N=264 (=B). Error calculated as mean distance of bootstrap predictions from the centroid predicted for each individual (see Fig. S4). A black dot denotes the predicted location of an individual and a colored dot indicates the ‘true’ location (colors proportional to measurement error in km).

Geo-located results for individuals passing a strict error filter demonstrated a dispersal distance for males approximately double that of females across all age classes (statistically significant only for the Y2-2.5 class due to low sample sizes; Fig. 6). This pattern was established as early as the Y1-1.5 group, indicating apparent male dispersal by that age. Smaller dispersal distances were found for fawns across both sexes (Fig. 6), again corroborating *a priori* biological expectations.

**Figure 6:**
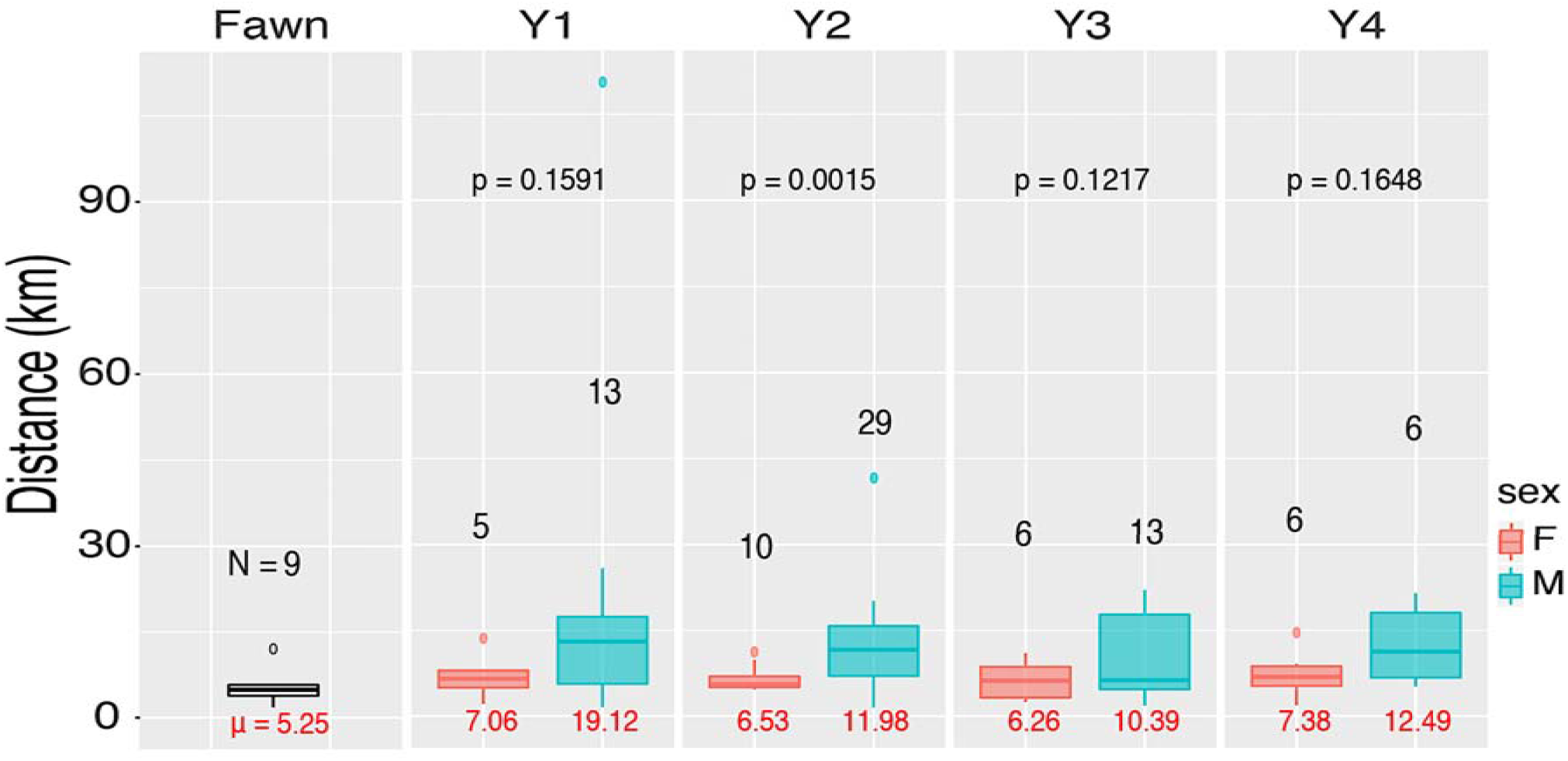
Inferred dispersal distances, partitioned by sex and age class for *N*=1,143 Arkansas white-tailed deer. Dispersal distances were calculated as the difference (in km) between the predicted and true locations, excluding all individuals with a mean prediction error of 10km (see Figs. 14-15). Sample sizes for each group are given in black, with the mean in red below each box plot. Within-age, p-values are reported for two-sample t-tests comparing means from males and females.

Patterns of genetic dissimilarity also showed an age x sex effect, with greater genetic dissimilarity for neighboring Y0-1 males than females, and with a shift towards reduced dissimilarity in males >5 (Fig. S5). This again supports the argument that male deer in Arkansas have dispersed by the Y1-1.5 age class, and potentially reflects age-biased reproduction. Males contributed disproportionately to their local gene pools by age 5 (i.e., producing offspring with resident females), thereby creating a pattern of lower genetic dissimilarity among neighboring individuals, regardless of distance.

## 4 DISCUSSION

### 4.1 Genetic footprints of historic management

The prolonged history of hunter-harvest, and subsequent long-range translocations into Arkansas and the surrounding region (Ellsworth et al., 1994; Holder, 1951), provide a sideboard to our study in that artificial long-distance movements such as these violate the assumptions inherent with many spatially-explicit methods. However, we found estimates of ancestry diversity and probability surfaces to be qualitatively robust in that they firmly recapitulated the record of historic translocations (Holder, 1951). This, in turn, necessitates that our results be placed within an historic context.

In the early 20th century, following many decades of over-hunting, the Arkansas deer population declined to <500 individuals (Holder, 1951). In response, Arkansas Game and Fish Commission (AGFC) implemented an extensive restocking program (1941-1951) involving as its basis three primary sources. The first was Howard County deer farm (southwestern Arkansas; Fig. 7), established from locally transplanted central Arkansas individuals (Wynn, 1943), and now located within the epicenter of population *k*1 (Figs. 2, 7). Its ancestry is shared elsewhere in the state, thus establishing it as an epicenter for local translocations.

**Figure 7:**
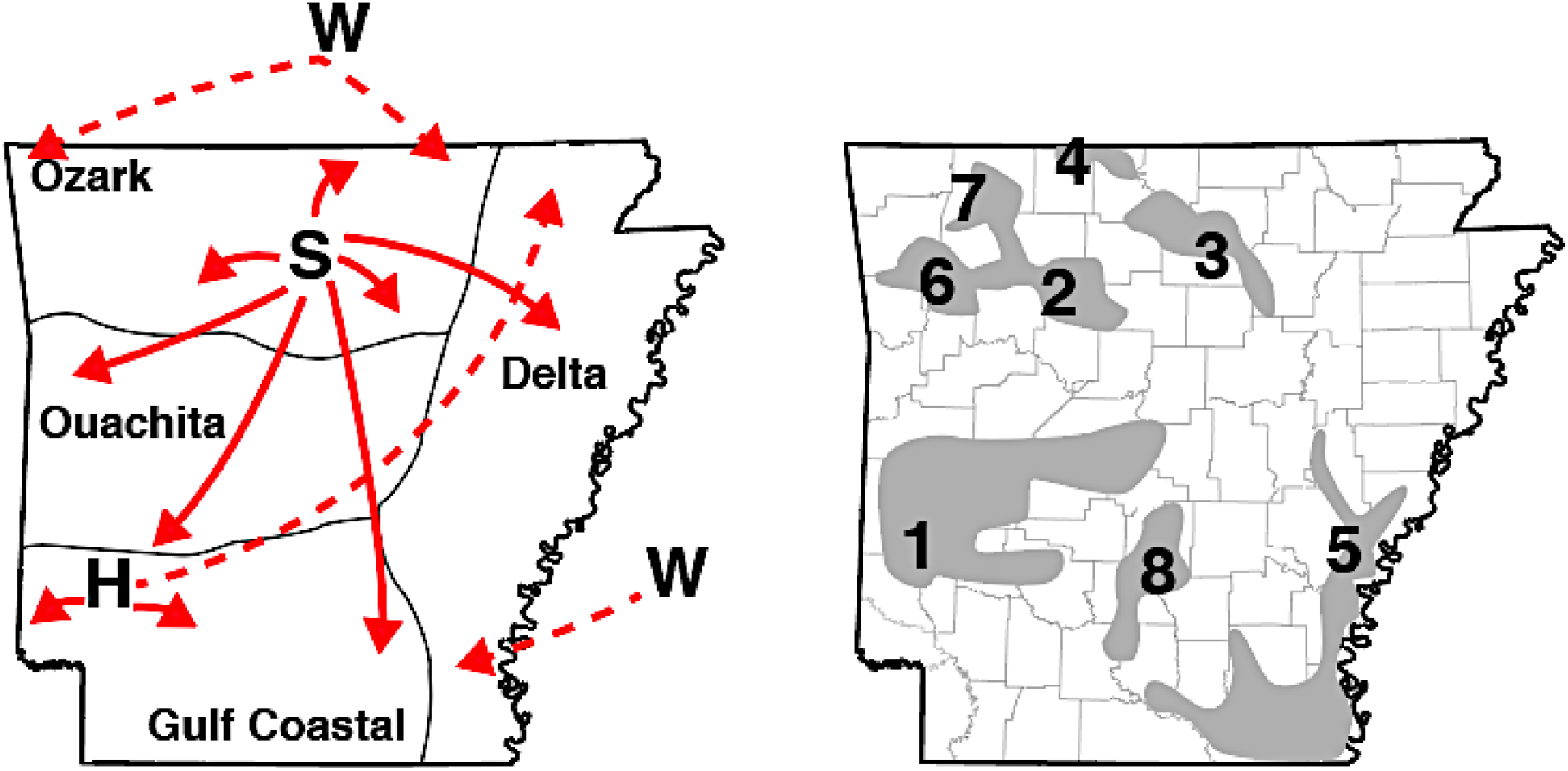
Proposed translocation pathways and refugial populations for Arkansas white-tailed deer, showing (left) translocation events supported by genetics along (dashed line), as well as those with anecdotal support given limited historical records (solid), from three primary stocking sources: Sandhill Game Farm in Wisconsin (W); ‘Sylamore District’ state refuge sites (S); and the Howard County Game Farm (H). Also shown are major inhabited regions of white-tailed deer (right) estimated from 1942-47 game surveys, excluding small regions (see full version in Online Supplementary Material). Putative inhabited regions are also annotated according to their hypothesized associations with contemporary genetic clusters (Figs. 1-2).

A second major source was the Sylamore District in the Ozark Mountains (Wood, 1944; Holder, 1951), where individuals were naturally abundant (Fig. S7). Individuals from this region have a consistently higher probability of assignment to *k*3 (Figs. 2, 7), as records indicate ∼81% of repopulated individuals in the Gulf Coastal Plain originated from there (Holder, 1951; Karlin, Heidt, & Sugg, 1989; Wood, 1944). Our analyses agree in that many individuals from southwestern Arkansas (Figs. 1, 6) reflect a mixed assignment to the ‘Sylamore’ cluster. They also comprised ∼36% of stocking efforts in the Mississippi Delta region (Karlin et al., 1989).

Our results also indicate some mixed assignment of individuals to *k*3, although a more widespread representation is found in *k*5. AGFC surveys (∼1942-1945) indicated deer as being relatively abundant in the southeastern Delta region (Figs. 7, S7), seemingly linked with re-population efforts following the disastrous 1927 Mississippi flood. This event, coupled with over-hunting, nearly extirpated deer in the region, save small numbers sustained by local sportsmen (Holder, 1951; Fig. S7). We hypothesize deer in the Delta region largely result from those efforts, given the contemporary homogeneity of ancestry assignments in this region.

A third (extraneous) source was the Sandhill Game Farm (Babcock, WI; Wood 1944). Records indicate that ∼64% of deer released into the Mississippi Delta region originated out-of-state, the majority from Wisconsin (Holder, 1951; Karlin et al., 1989). Our Wisconsin samples were unanimously assigned to a gene pool prominently represented in the southern Delta region (*k*8; Fig. 2), firmly establishing its genetic legacy as extending from imported deer. Mitochondrial haplotypes putatively originating from Wisconsin were also uncovered in Missouri, Kentucky, and Mississippi (Budd, Berkman, Anderson, Koppelman, & Eggert, 2018; DeYoung et al., 2003; Doerner et al., 2005).

All genetic ‘clusters’ that lacked spatial cohesion in our analyses can be connected to the three major stocking sources involved in earlier restoration efforts (per historic records). The remaining sub-populations, primarily in the Ozarks (*k*2, 4, 6, and 7) may represent natural re-colonization from refugial populations (Figs. 2, 7), an hypothesis supported by early census data (Figs. S7, S8). If this was not recognized, then any analysis of deer genetic structure across Arkansas would instead infer unusually high rates of gene flow among these geographically distant regions. This, in turn, would preclude any analysis aimed at linking variation in genetic structure to environmental factors.

### 4.2 Contemporary genetic structure

A primary focus in our study was the contemporary genetic structure of Arkansas deer, especially regarding how these data promote an understanding of landscape resistance to dispersal (Hemming-Schroeder, Lo, Salazar, Puente, & Yan, 2018; Kelly et al., 2014). Yet several prerequisites are apparent in this regard. For example, one apparent question previously constrained by technological limitations is the degree to which potential patterns of genetic variability in deer have been conflated by anthropogenically-mediated translocations.

We addressed this issue by utilizing next-generation molecular techniques to derive highly variable markers across the genome. We also implemented advanced geospatial procedures that visualized the spatial transitions inherent within deer ancestry in such a way that translocation histories were not a limiting factor. We did so by interpolating our assignment probabilities from ADMIXTURE, then applying Simpson’s diversity index as a means of reducing the probabilities to a single vector. We also polarized stochastic *versus* deterministic processes by examining locus-specific patterns within our transitions.

Previous studies concluded that inferences from landscape genetic analyses were limited at best, due to complex interactions between historic translocations and subsequent population growth (Leberg et al. 1994; Leberg and Ellsworth 1999; Budd et al. 2018). However, many of these studies relied upon legacy molecular markers (e.g., mtDNA or reduced panels of microsatellite DNA markers) that capture substantially less polymorphism than do next-generation methods (Hodel et al., 2017; Jeffries et al., 2016; Lemopoulos et al., 2019). Recent studies at more refined spatial scales have supported large-scale geographic barriers (e.g. rivers, highways) as being semipermeable to gene flow (Kelly et al., 2014; Locher et al., 2015; Miller et al., 2020; Robinson et al., 2012). This, despite the potential complication caused by re-stocking efforts, and in accordance with radio-telemetry data (Peterson et al., 2017).

Several scale-dependent issues may modulate the degree to which translocations obfuscate landscape genetic patterns. First, as spatial scale increases, so does the probability of sampling individuals whose genetic dissimilarity reflects translocation rather than natural gene flow via dispersal. Second, the most obvious environmental effects that drive deer dispersal have the greatest probability of being recovered in an analysis, even if they represent spurious signals driven by translocation. However, landscape-level analyses of dispersal can still be informative, despite those rapid transitions between historic and contemporary conditions (Epps & Keyghobadi, 2015) that may obfuscate relationships. For example, large-scale environmental features also drive spatial genetic patterns in invasive species, despite reduced levels of genetic diversity (Lopez, Hurwood, Dryden, & Fuller, 2014; Sacks, Brazeal, & Lewis, 2016; Zalewski, Piertney, Zalewska, & Lambin, 2009). Such patterns are also rapidly manifested within populations occupying habitats recently modified by anthropogenic developments, such as large urban centers (Beninde et al., 2016; Combs, Puckett, Richardson, Mims, & Munshi-South, 2018; Kimmig et al., 2020).

The analytical artefacts derived from translocation are expected to be not only scale-dependent but also a function of specific analytical assumptions. For example, classic tests of ‘isolation-by-distance’(IBD) implicitly assume a negative relationship between geographic proximity and pairwise patterns of genetic distance (Meirmans, 2012; Rousset, 1997). Yet, translocations are demonstrably inconsistent with this expectation. Introduced individuals reflect levels of genetic similarity that are demonstrably inconsistent with spatial and/ or environmental distance (i.e., ‘resistance’) that now separates them. This, in turn, incorporates spurious signals within the landscape inferences subsequently derived.

The assignment method used in our study yielded patterns of genetic similarity that reflected translocations as depicted in historic records (Fig. 7). We sought to more appropriately expose landscape features that could potentially modulate deer dispersal by effectively ‘removing’ this artificial signal. Our approach (i.e., reducing the dimensionality of assignment probabilities) produced relatively straightforward predictions about the ensuing metric: Areas with uninterrupted movement of individuals should have increased homogeneity regarding interpolated ancestry assignments. By the same logic, regions representing barriers to movement among regions should likewise demarcate transitions in interpolated ancestries. We acknowledge that spatial assignments can also be vulnerable to artefacts, such as an over-fitting of discrete clusters within otherwise continuous populations (Bradburd et al., 2018). Yet despite this, they are demonstrably effective at identifying linear barriers to gene flow (Blair et al., 2012). Coincidentally, our approach using summarized ancestries revealed numerous linear subdivisions that aligned with major landscape barriers, such as rivers in Arkansas (Fig. 3). Other recognized transitions approximately corresponded with large urban centers. Our argument is that approaches involving an assumption of stepwise gene flow (implicit with pairwise genetic distances) may generate conclusions that either lack the necessary nuances or are altogether incorrect. This is apparent when the ‘surfaces’ inferred using Simpson’s diversity of ancestry assignments (Fig. 3) are contrasted with EEMS results (Fig. S1).

### 4.3 Separating stochastic *versus* deterministic processes in clinal patterns

When deriving clinal patterns, it becomes difficult to disentangle contemporary demographic processes from analytical idiosyncrasies. As a means of clarification, we generated multi-locus genomic patterns under the expectation that the relative importance of stochastic *versus* deterministic processes could be derived from the manner by which genome-wide averages deviated from locus-wise clines (Barton, 1983; Barton & Hewitt, 1985; Slatkin, 1973; Vasemägi, 2006). We found eight transects that varied either in the steepness of genetic transitions (β), or the directionality of allelic frequency change (α). Interestingly, ‘α-dominant’ transects were found within the Ozark region, previously hypothesized as being naturally re-colonized from local refugia (i.e., *k*2 x *k*3E/S, *k*2 x *k*6, and *k*4 x *k*7; Figs. 4, S2).

Under this hypothesis, genetic drift along the leading edge of population expansions would generate two genetic reverberations: The random, directional fixation of alleles, and an apparent inflation of population structure. Both were observed in our data (Fig. 1, 4). The hypothesis is also supported by state-wide surveys (1940’s) that found deer in habitat associated with each Ozark sub-population during, and immediately following, the low ebb of deer in Arkansas (Figs. 7, S8). The transects likewise displayed near-zero β values, suggesting the absence of a ‘true’ barrier to dispersal. Here, the expectation would be that a hard barrier between discrete populations would reflect changes in allele frequencies around an inflection point.

One interpretation is that genetic drift at the edge of an expanding refugium creates a ‘rolling’ founder effect, with alleles either over-(α>0) or under-represented (α<0) across the re-colonized territory (Excoffier & Ray, 2008; Hallatschek & Nelson, 2008). This phenomenon— termed ‘gene surfing’—can generate wholly neutral patterns that appear to be adaptive (Peischl, Dupanloup, Bosshard, & Excoffier, 2016; Travis et al., 2007), and may explain the presence of numerous α-outlier loci in those transects (Fig. 4).

Interestingly, all our β-dominant transects crossed major rivers: Transect *k*3 x *k*6 spanned the Arkansas River; *k*3 x *k*5 the White River; and *k*1 x *k*8 the Ouachita River. Transect *k*3 x *k*4 crosses the smaller and shallower Buffalo River, which may explain the depressed variability in cline steepness (Fig. 4). The coincidence of these patterns with major rivers suggests barriers to individual movement, in that non-zero β values implicate variability in the ‘steepness’ of clinal transitions around an inflection point [i.e., with rates of change in allele frequencies either steep (β>0) or wide (β<0)].

Classically, clinal patterns are established either by selection or hard boundaries to gene flow (Endler, 1973; Slatkin, 1973), but our analyses (Fig. 4) revealed little evidence of selection-driven outliers. However, spatially variable gene flow, when coupled with drift, can yield clinal patterns across a considerable proportion of the genome without necessitating selection (Vasemägi, 2006). Selection could feasibly be involved, at least for regions where different deer subspecies were translocated, in that subspecific crosses have elicited fitness impacts such as dystocia (abnomal maternal labor due to shape, size, or position of the fetus; Galindo-Leal and Weber 1994). The genetic costs of inter-subspecific stocking are recognized (Hopken, Lum, Meyers, & Piaggio, 2015), and manifested as an anomalous variation in breeding time among other southern-recovered deer populations (Sumners et al., 2015). Some limited evidence also for reproductive isolation among mule deer subspecies (*Odocoileus hemionus*) as driven by pheromonal differences (Müller-Schwarze & Müller-Schwarze, 1975).

### 4.4 Management implications

Our results identified a diagnosable genetic signature of historic translocations within the genomes of extant Arkansas deer populations. This simultaneously underscores the success of early restocking efforts, while also reiterating long-standing concerns about the genetic and/ or phenotypic impacts of anthropogenic translocations (Meffe & Vrijenhoek, 1988). It also reinforces the need to formalize as an explicit management consideration the intraspecific taxonomy of deer, both to better understand the (yet to be seen) evolutionary consequences of past translocations, and to provide a more proactive baseline going forward (Cronin, 2003; Gippoliti, Cotterill, Groves, & Zinner, 2018).

Resource agencies, in particular, should prioritize those populations that retain endemic genetic diversity, and by so doing, acknowledge the importance of preserving evolutionary legacy (Crandall, Bininda-Emonds, Mace, & Wayne, 2000). Such recognition would capitalize on the many years of natural selection that have operated on those populations prior to anthropogenic interference, which may have generated adaptive genetic variation necessary to withstand changing conditions, or mediated an evolutionary response to disease (e.g., in the *PRNP* gene; Chafin et al., 2020). This further emphasizes the importance of integrating genomic methods for heavily managed species (Flanagan et al., 2017), given their recognized lack of intraspecific adaptive variation [i.e., a ‘Darwinian shortfall’ in biodiversity conservation (Diniz-Filho, Loyola, Raia, Mooers, & Bini, 2013)].

It is noteworthy that the current boundaries for the AGFC ‘Deer Management Units’ (DMUs) (Meeker et al., 2019) are remarkably consistent with many of the population boundaries identified in our study. This validates the continued use of these ecosystem-based management units for the application of locally-appropriate management and harvest regimes.

Another valuable application of our results would be disease containment, particularly given that CWD is a significant challenge globally for the management of cervids (Rivera, Brandt, Novakofski, & Mateus-Pinilla, 2019; Uehlinger, Johnston, Bollinger, & Waldner, 2016). The spread of CWD will be more rapid within rather than among populations, and the cumulative risk of disease spread to new populations will depend on both dispersal and disease prevalence. Thus, potential mitigation efforts should work to constrain disease transmission within CWD-affected areas as well as limiting the outward dispersal of individuals from those areas. One approach to reduce dispersal is to reduce the density of yearling males, which often comprise >50% of emigrating individuals (McCoy, Hewitt, & Bryant, 2005; Nelson, 1993; Nixon, Hansen, Brewer, & Chelsvig, 1991). Our patterns of age- and sex-biased dispersal distances (Fig. 5, 6) largely recapitulate these expectations at a local level. Removal of young males to slow dispersal also may be compatible with the application of male-focused harvest strategies predicted to reduce disease prevalence (Potapov, Merrill, Pybus, & Lewis 2016). A similar action would be to reduce local population densities that serve to drive dispersal, in that robust population numbers elevate social pressure as well as resource competition (Long et al., 2008; Lutz et al., 2015; Shaw, Lancia, Conner, & Rosenberry, 2006). Other harvest strategies have been implemented that effectively decrease emigration while focusing on meeting specific demography goals including: actively decreasing overall population density, elevating the proportion of mature males in the population, and increasing the female component of the sex ratio (Brothers and Ray, 1975; Hamilton et al., 1995; Shaw et al., 2006; Long et al., 2008; McCoy et al., 2005; Lutz et al., 2015). However, mature male deer tend to demonstrate higher CWD prevalence than other demographic groups, and management strategies that maximize this component of the population may exacerbate disease dynamics (Miller, Runge, Holland, & Eckert 2020). Additionally, our findings indicate that disease surveillance strategies should include efforts that focus on detecting ‘breaches’ between genetically distinct regions (those bordered by major rivers and urban centers, as herein; Fig. 3) to further direct management and response efforts.

### 4.5 Utility of genomic approaches for white-tailed deer management

Beyond providing a nuanced view of historical management processes, the genomic methods used herein offer several improvements beyond ‘legacy’ approaches. For example, they provide vastly increased statistical power for individual-level genetic analyses, such as estimating relatedness (Lemopoulos et al., 2019), or parentage assignment (Flanagan & Jones, 2018). This is also accomplished at a fraction of the cost-per-marker (Puckett, 2017). Evidence for this is found in our geolocation results (Fig. 5, S3-S4), where a lower dispersion of predicted natal locations were achieved when compared with previous attempts using microsatellites (Green et al., 2014). The genome is also more densely sampled and these data can be used in conjunction with the analytical methods used herein to infer both environmental associations and adaptive genetic variation (Martin et al., 2020), or detect anthropogenically-mediated change through time (Chafin et al., 2019).

SNP-based methods have a particular advantage with regards to scalability (i.e., such as in a multi-state consortium of wildlife management agencies). Large datasets (as herein) may allow a subset of highly consistent, maximally informative markers to be extracted that are broadly applicable across states and regions. In ddRAD (and other RADseq methods), much sequencing effort may be wasted on low-occupancy or systematically-missing loci, a result of variability introduced during: Library preparation (Escudero et al., 2014); Bioinformatic processing (Eaton, 2014); Mutation-disruption (Rubin et al., 2012), or; Insufficient sequencing coverage (Huang & Knowles, 2016). Additionally, many loci may be monomorphic, contain low-frequency mutations, or be uninformative at the scale examined. Methods to optimize cost-efficiency can be applied to consistently assay target loci (e.g., Chafin et al., 2018) by using targeted-enrichment approaches [e.g., Rapture (Hoffberg et al., 2016)] or those amplicon-based [GT-Seq (Campbell, Harmon, & Narum, 2014)], thus reducing the total costs to <$10 per sample.

Selecting an appropriate approach depends upon desired throughput capacity, yet all share the need for an initial database of genetic variation from which to assays can be generated. For white-tailed deer, our approach would first be applied to generate a broad geographic reference database (i.e., at a national or multi-regional scale), followed by assay design for a *post hoc* method (such as that of Campbell, Harmon, & Narum, 2014). This would allow our methodology to be deployed so as to generate a sampling sufficiently dense for management-oriented population genetic analyses such as natal geolocation. This, in turn, would facilitate the detection of illegal carcasses, deer trafficking, migrant detection, or characterization of historic translocation (as herein). Moreover, such an approach would be fully replicable across laboratories and agencies, thereby facilitating inter-agency and multi-state collaboration.

Although dense spatial and genomic sampling may offer a composite resolution (e.g., Chafin et al., 2020; Linck et al., 2019), it does not necessarily supplant other approaches. For example, sampling density was shown herein to be a particular limiting factor in geolocation analyses, with high prediction error in regions of the state sparsely sampled (Fig. 5). Although we did not directly compare our methodology with that based on a microsatellite dataset, we suspect greater error in the latter given decreased information content associated with the much reduced dataset. Genetic input from unsampled populations (e.g., long-distance migrants from out-of-state) found at edges of our sampling areas could also inflate assignment uncertainty (see Fig. 3), as could extremely uneven sampling (Lawson, van Dorp, & Falush, 2018). We are also unsure of the degree to which long-distance migrants could influence our clinal analyses (e.g., via a ‘smoothing’ effect; Smith and Weissman 2020), nor fluctuation that may occur over brief time scales (as observed in other study systems; Chafin et al., 2019). Thus, we caution that the deployment of these methods at multi-state or regional scales should focus on dense, spatially expansive sampling that employs both spatial and temporal replicates, where possible (Short Bull et al., 2011). Both priorities are facilitated via the aforementioned enrichment methods that drastically reduce per-sample cost-efficiency.

## ACKNOWLEDGMENTS

The authors thank hunters and wildlife managers statewide who collected WTD tissue samples. We give particular thanks to A.J. Riggs for helping to coordinate sample collection. We also thank L. Graham for contributions to study design, logistics, and DNA extraction, S.M. Mussmann for assistance with geospatial analyses, and J.F. Pummill for facilitating access to computational resources. These were provided by the Arkansas High Performance Computing Center via multiple National Science Foundation grants, the Arkansas Economic Development Commission, as well as an NSF-XSEDE Research Allocation (TG-BIO160065) that provided access to the JetStream cloud service. This research project was funded by the State of Arkansas and the Wildlife and Sport Fish Restoration Program (#AR-W-F15AF00896), and also in part by the Pittman-Robertson Wildlife Restoration Grant Program (#AR-W-F18AF00020) of the U.S. Fish and Wildlife Service through an agreement with the Arkansas Game and Fish Commission. Additional funding was also provided by several generous endowments from the University of Arkansas: Distinguished Doctoral Fellowships (TKC, ZDZ), the Bruker Professorship in Life Sciences (MRD), and the Twenty-First Century Chair in Global Change Biology (MED). The opinions expressed herein do not necessarily reflect the views or policies of the Arkansas Game and Fish Commission or the U.S. Fish & Wildlife Service. In addition, product references do not imply endorsements.

## AUTHOR CONTRIBUTIONS

All authors contributed to study design and conceptualization. MCG, CRM, and JRB coordinated sample collection and curation of samples; TKC, BTM, ZDZ, and MRD contributed to molecular work; TKC, ZDZ, MRD, and MED planned the analysis approach; TKC and BTM wrote code for analysis and contributed exploratory analyses; TKC performed bioinformatic work, assembled data, and performed analyses; TKC and ZDZ prepared figures; TKC and ZDZ drafted the manuscript and all authors revised the manuscript and contributed to the final version.

## DATA AVAILABILITY STATEMENT

Raw sequences are accessioned in the NCBI Sequence Read Archive (SRA) under BioProject PRJNA690954; Assembled SNP data and sample metadata is available via the Open Science Framework (doi: 10.17605/OSF.IO/T82RV); codes and custom scripts developed in support of this work are also available as open-source via GitHub under the GNU Public License: github.com/tkchafin (and as cited in-text).

## Supplementary Figures and Tables

**Table S1:**
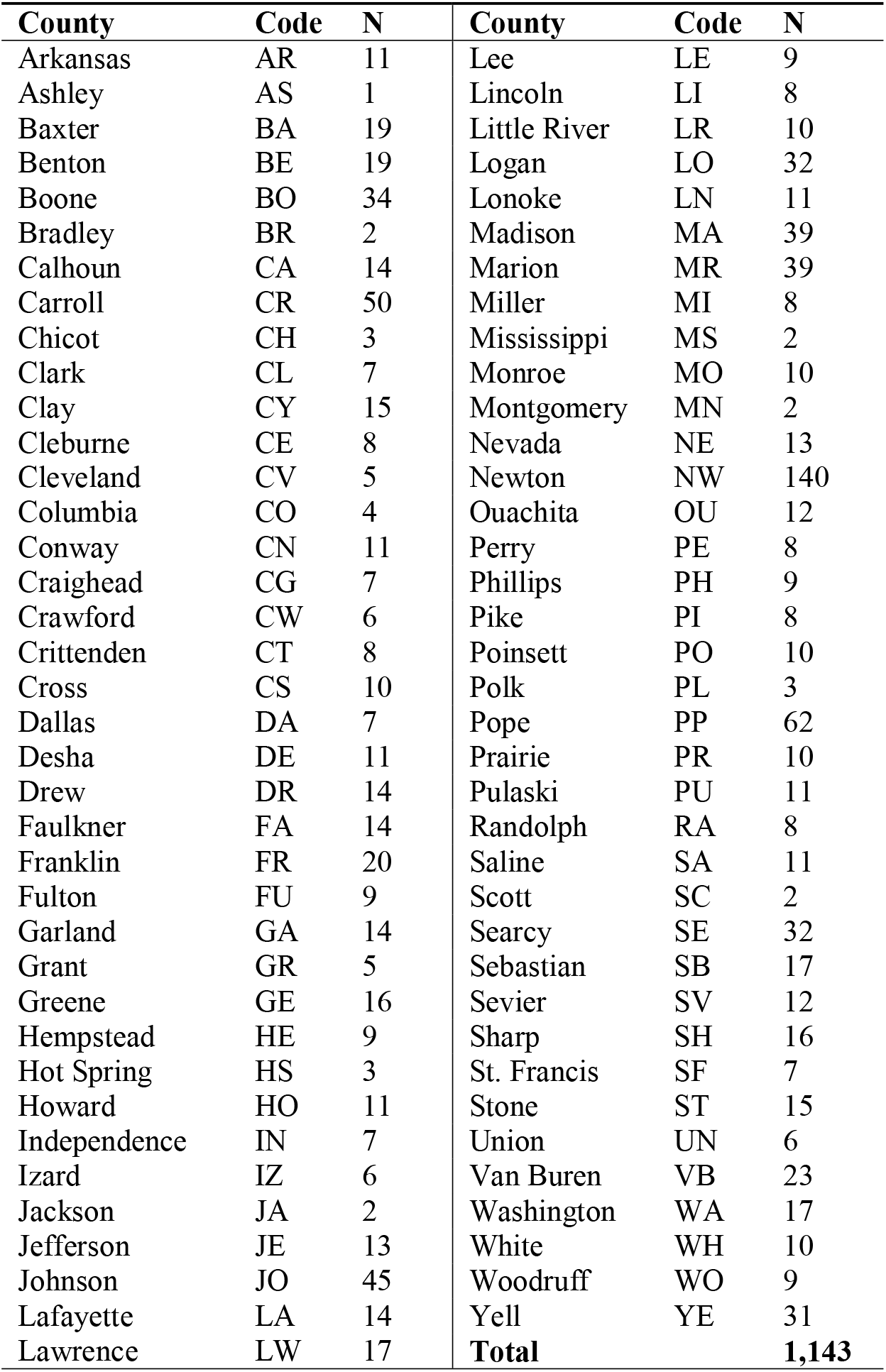
Numbers (=N) of Arkansas white-tailed deer genotyped across ∼35,000 SNP loci. Samples were collected from 75 counties (=County) from 2016-2019. Code indicates standard 2-letter county abbreviation. Samples removed due to missing data are not included.

**Figure S1:**
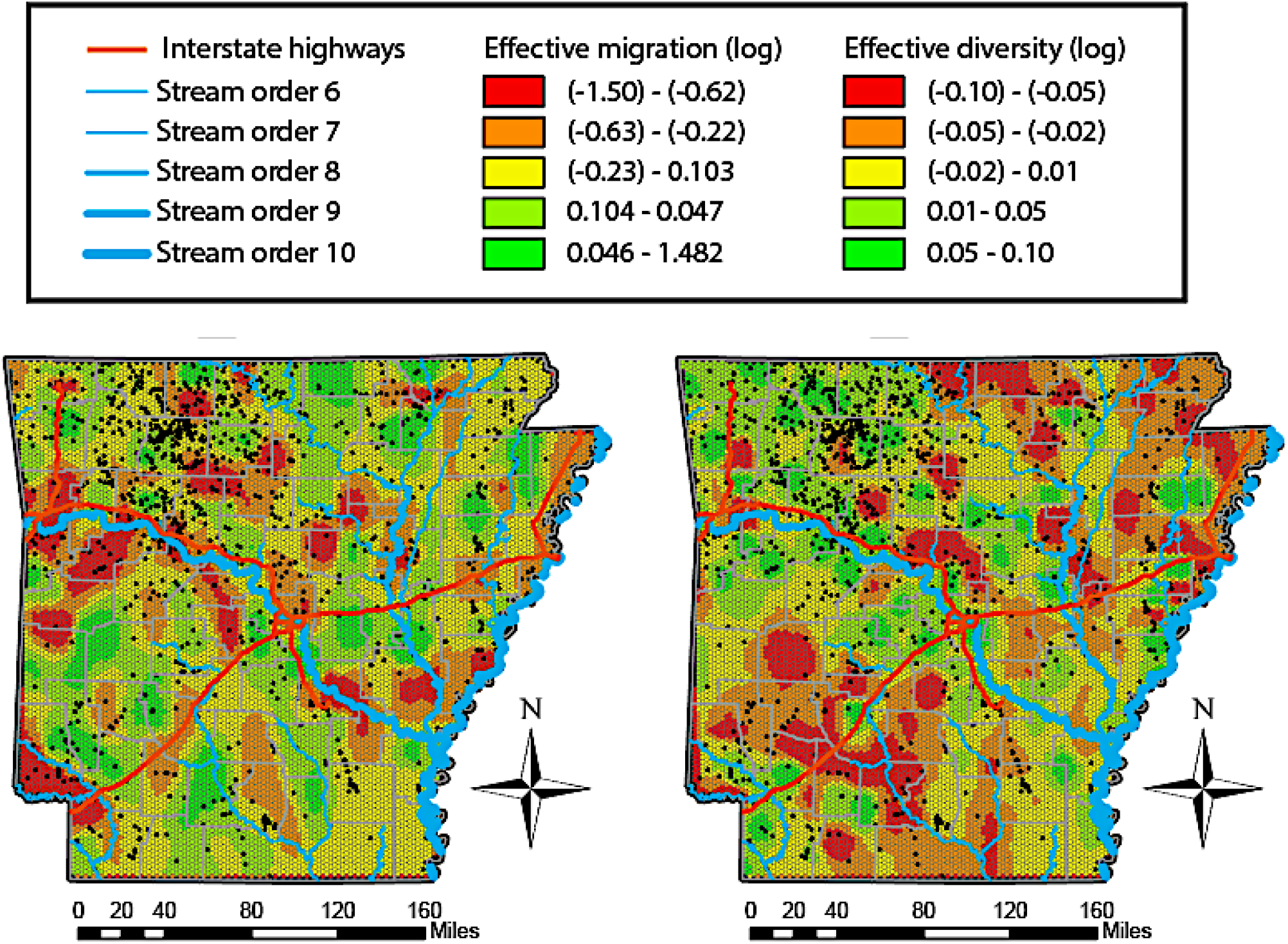
Effective migration rates and intra-population diversity. (log_10_ scale) for Arkansas white-tailed deer, calculated from effective migration surfaces (EEMS). Rates are plotted according to colored bin, with divisions calculated as natural breaks using the Jenks algorithm in ArcMAP.

**Figure S2:**
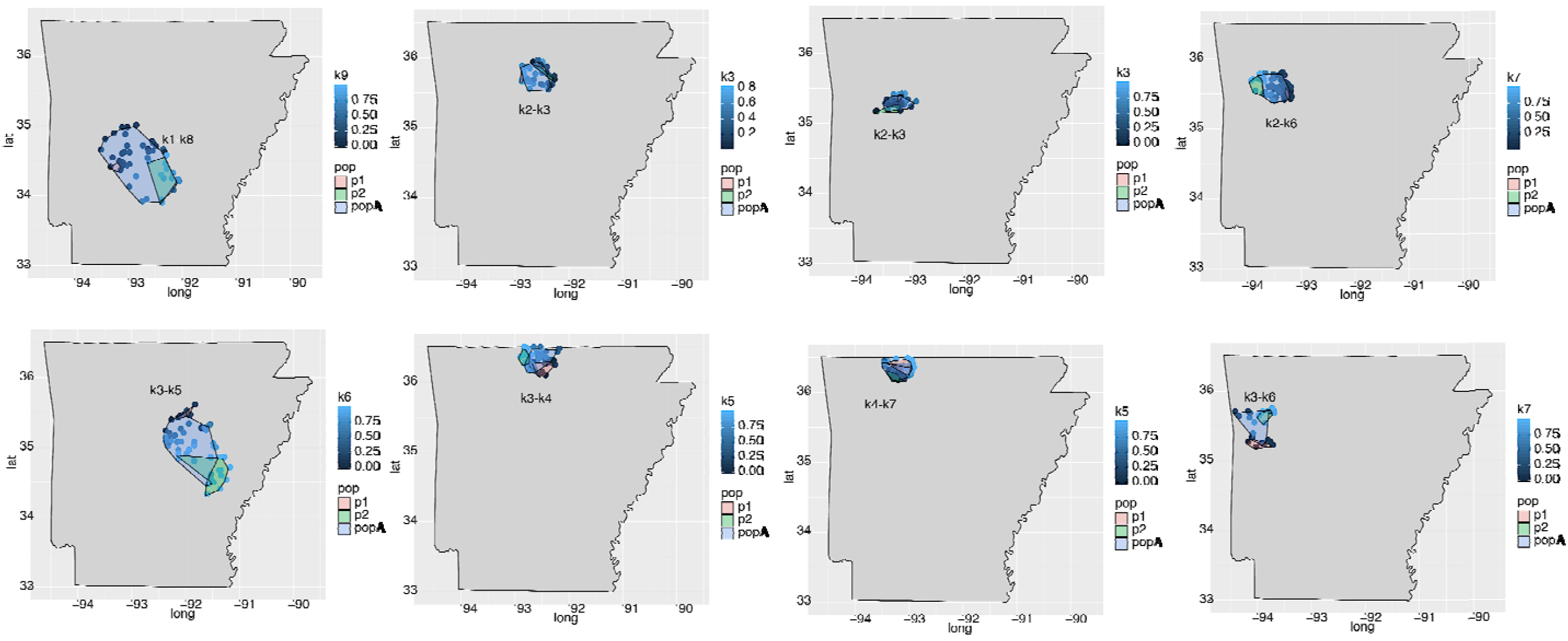
Sampled transects employed for genomic cline analysis of Arkansas white-tailed deer, with selected groups within each coalesced within colored hulls: Reference population 1 (p1; red); Reference population 2 (p2; green); and putative admixed individuals (popA; blue).

**Figure S3:**
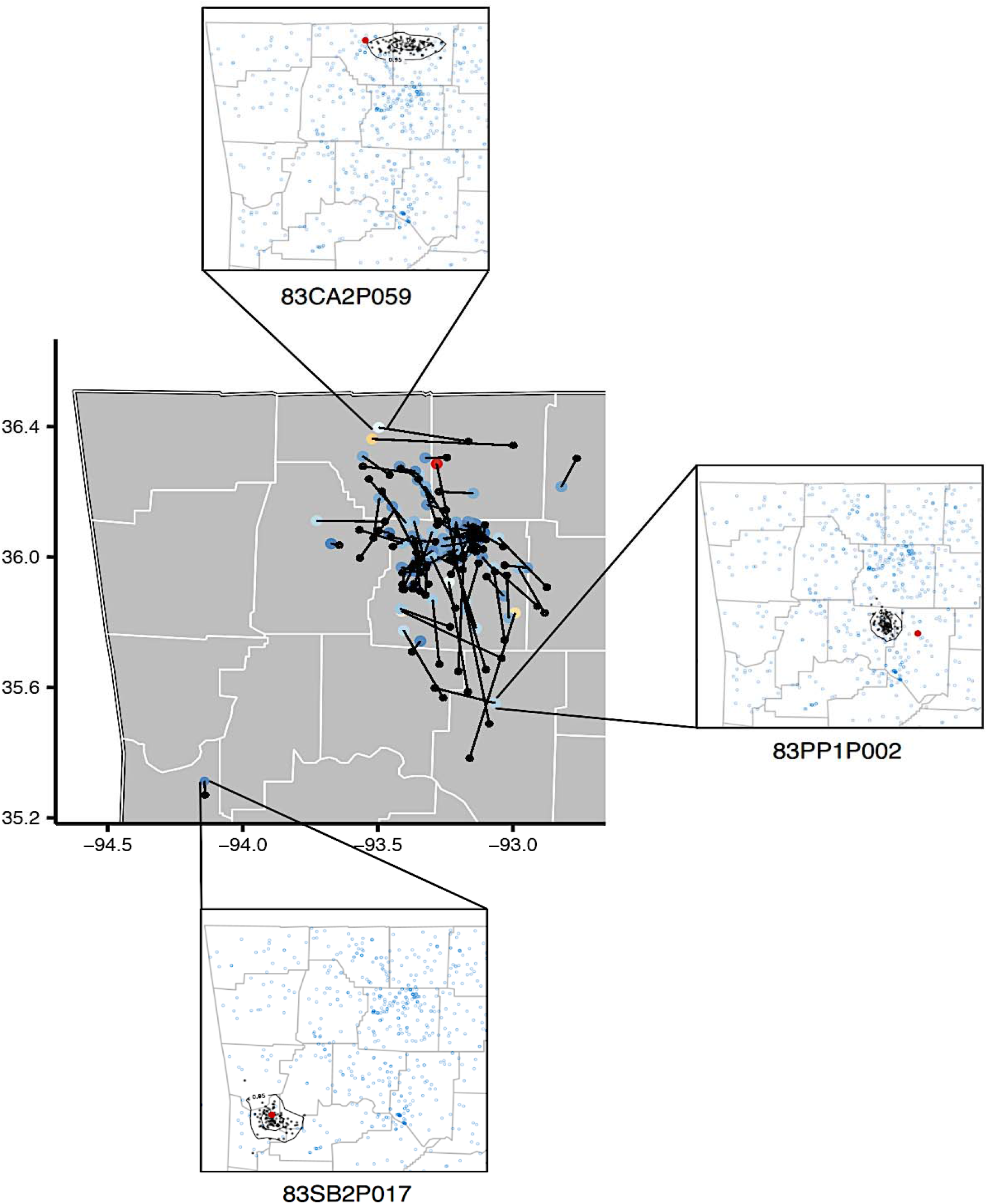
LOCATOR Geo-located results for CWD-positive Arkansas white-tailed deer derived via LOCATOR. Individuals are represented by dots, pairs of which are connected by a single line. A black dot denotes predicted location, while a colored dot indicates ‘true’ (observed = sampled) location (color proportional to distances separating each). Inserted figures offer full prediction results for three selected samples, with bootstrap estimates demarked by a 95% contour.

**Figure S4:**
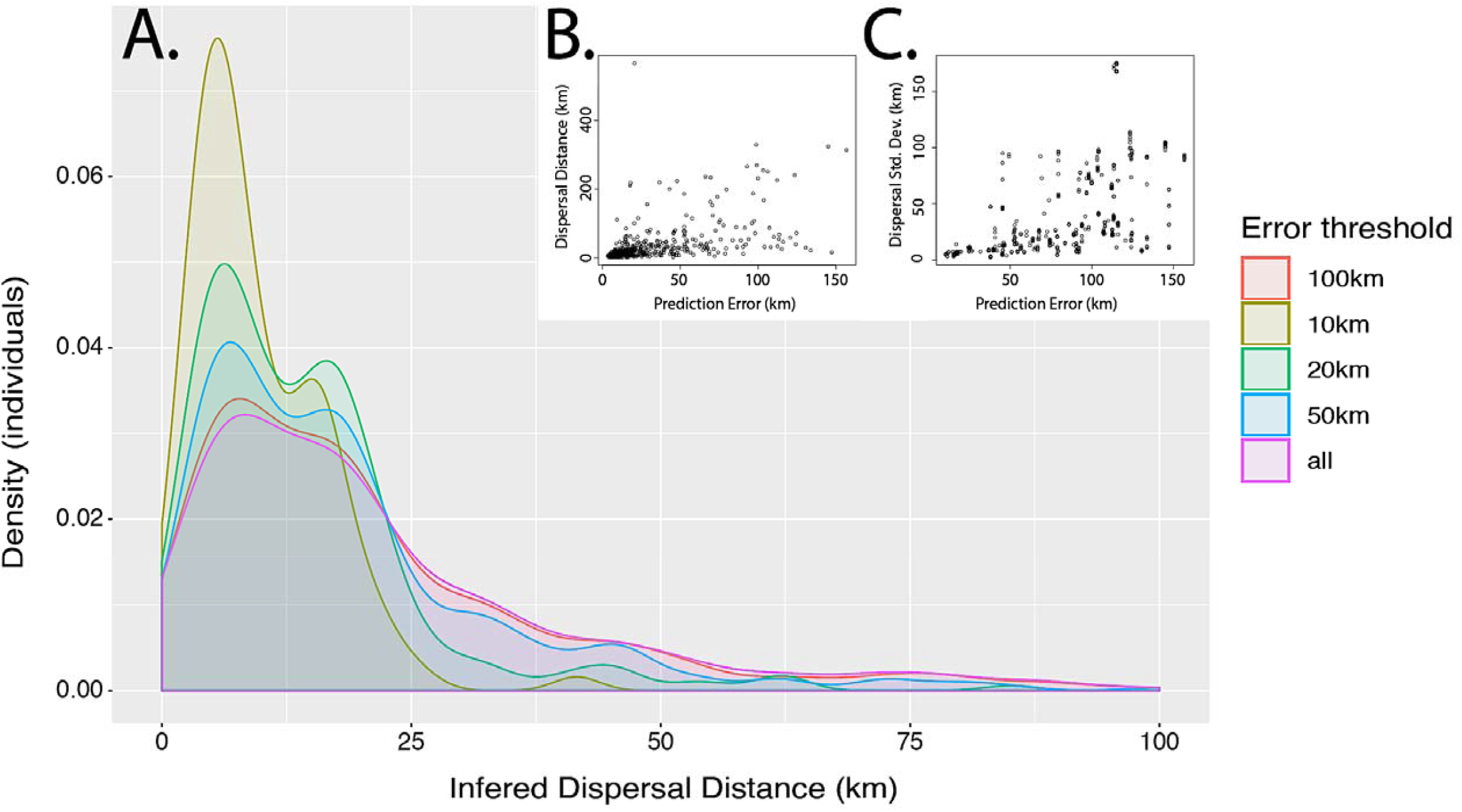
Effects of error thresholds on inferred dispersal distances for Arkansas white-tailed deer, as predicted by LOCATOR analysis. Inferred dispersal kernels were computed at several error thresholds: (A) Error computed as mean distance of predicted coordinates for each individual at 100 bootstrap estimates from predicted centroid (=interpretation of localized vs. dispersed predictions). Dispersal distances calculated as difference (in km) between predicted centroid versus ‘true’ (observed) location. Inferred dispersal distances generally increase for individuals with larger prediction error; (B) Larger stochastic variation in centroid location for individuals with lower predicted precision among bootstraps; (C) Standard deviation of dispersal distances for the latter computed in a sliding window of 10km along the x-axis (prediction error).

**Figure S5:**
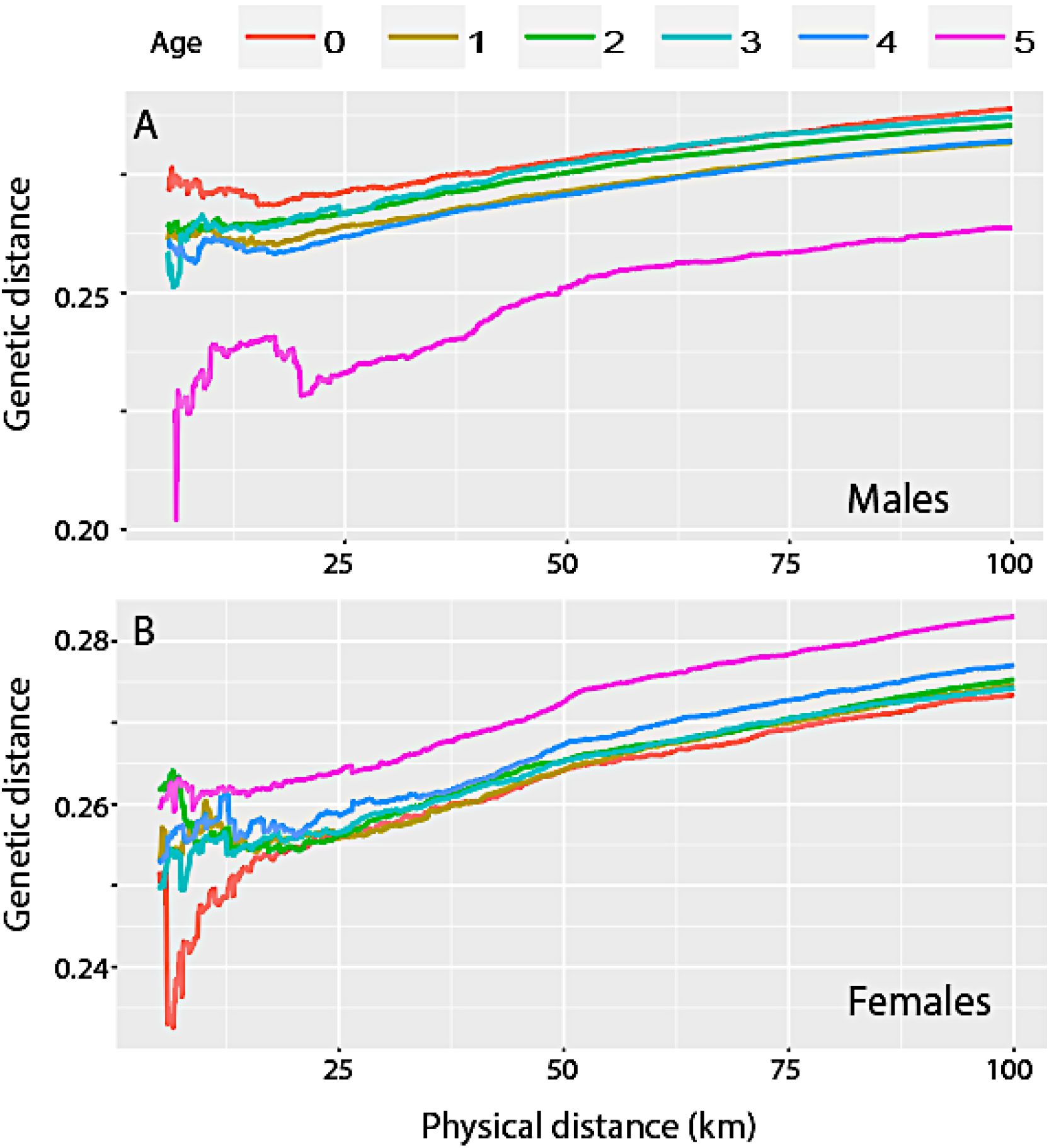
Spatial patterns of genetic dissimilarity among Arkansas white-tailed deer partitioned by age and sex. Genetic distances between individuals and neighbors derived from 5,000 randomly sampled SNPs depicted across physical distance (x-axis). Results depict different cohorts of males (A) and females (B) by age.

**Figure S6:**
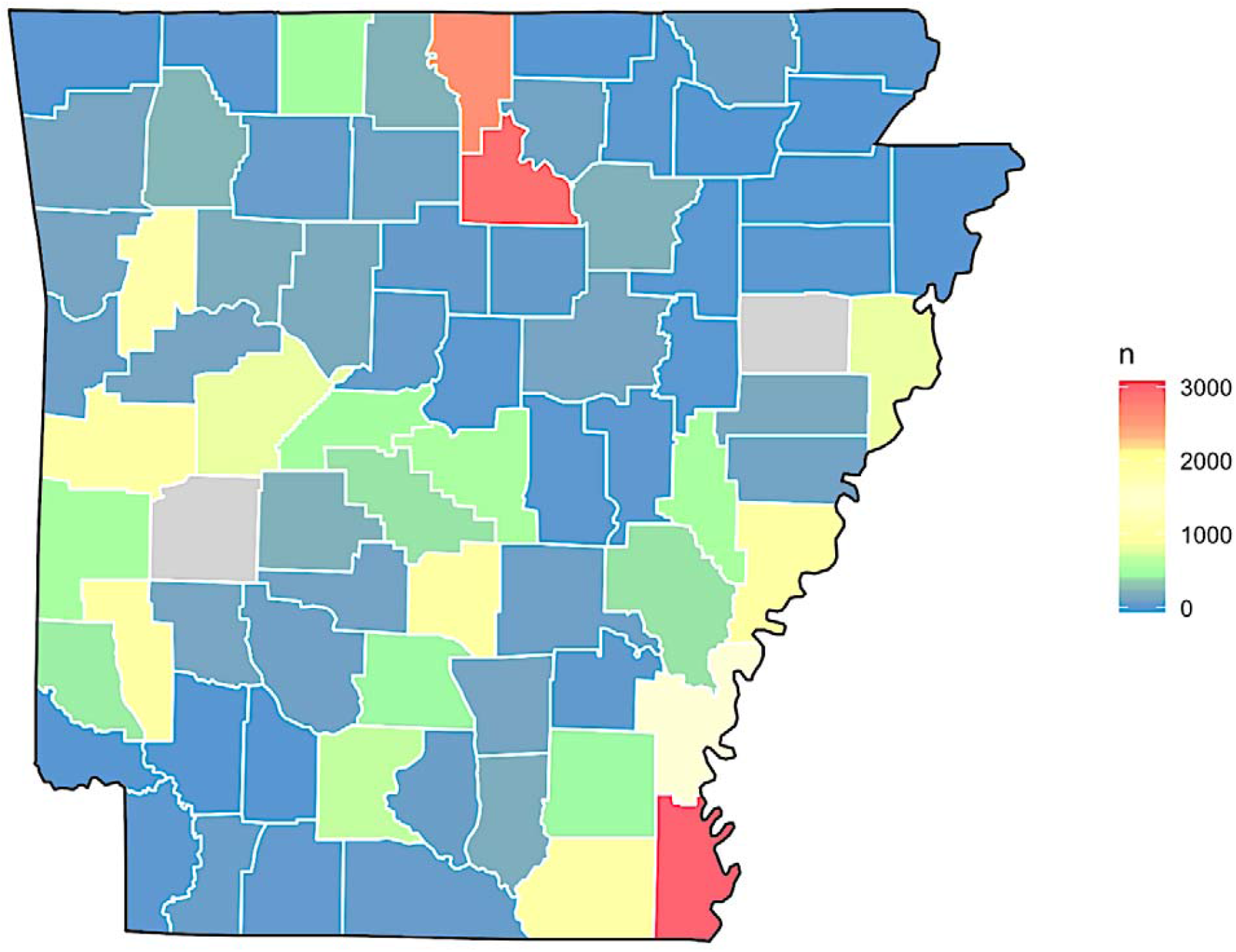
Census estimates for Arkansas white-tailed deer from surveys in 1942-1946.

**Figure S7:**
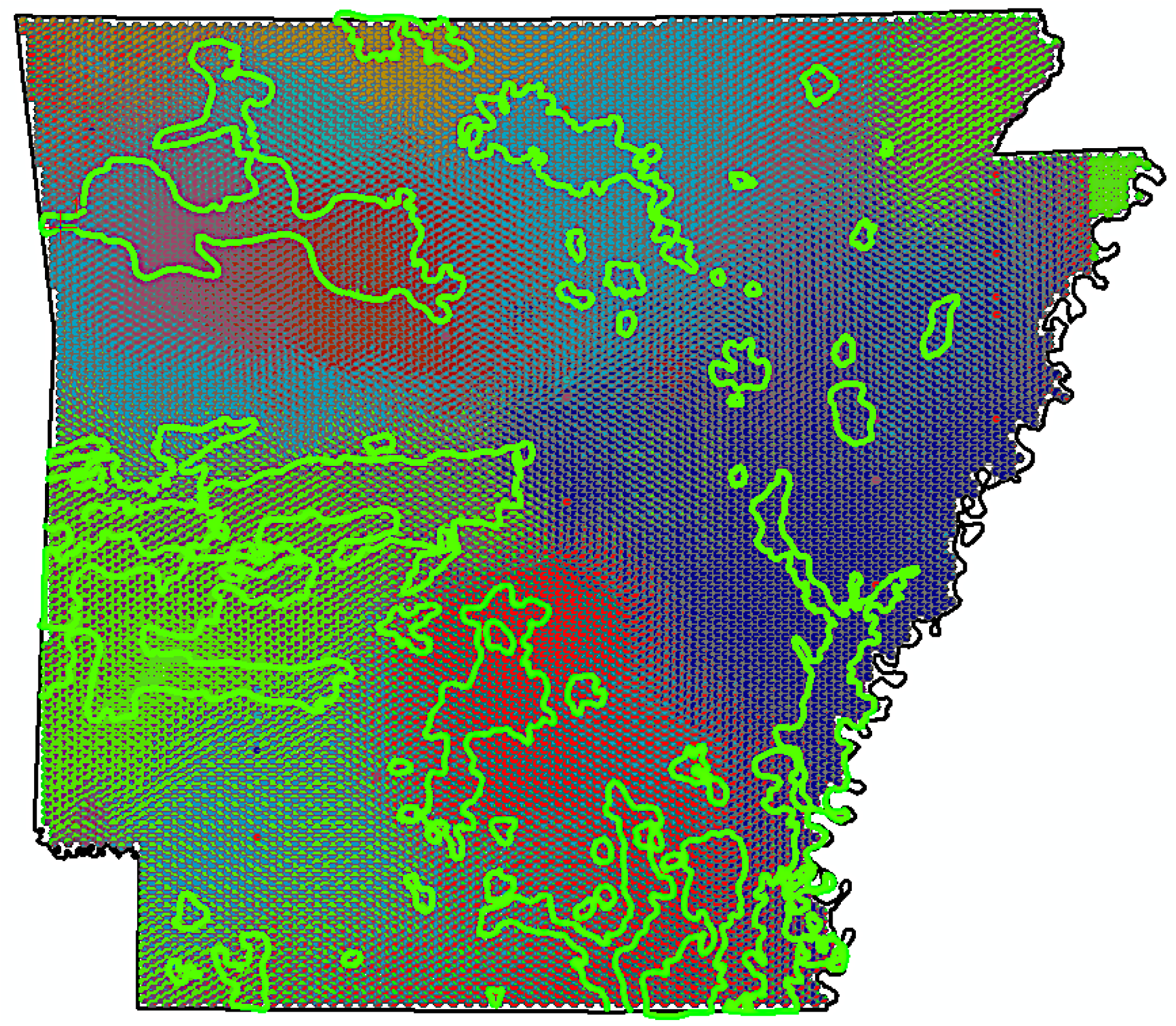
Subpopulation probabilities overlain with ‘deer occupied territories’ in Arkansa circa 1942-1947. (in green). Polygons representing deer occupation were manually compiled from the Wildlife and Cover Map of Arkansas (1942-47), prepared by Arkansas Game and Fish Commission in cooperation with U. S. Fish and Wildlife Service as one aspect of a Federal Aid Project (drawn by Flaun M. Tolar). Map accessed from the University of Arkansas Library Arkansas Collection (Special Collections). A higher resolution geo-referenced version available at a later date (reproduction/ copy permissions granted by University of Arkansas Libraries).

## Notes

### Competing Interest Statement

The authors have declared no competing interest.

### Summary of Updates

Minor changes in discussion and acknowledgements

